# A comparison of gene expression and DNA methylation patterns across tissues and species

**DOI:** 10.1101/487413

**Authors:** Lauren E. Blake, Julien Roux, Irene Hernando-Herraez, Nicholas E. Banovich, Raquel Garcia Perez, Chiaowen Joyce Hsiao, Ittai Eres, Claudia Chavarria, Tomas Marques-Bonet, Yoav Gilad

**Affiliations:** University of Chicago, Department of Human Genetics, Chicago, IL.; Department of Biomedicine, University of Basel, Basel, Switzerland.; Swiss Institute of Bioinformatics, Lausanne, Switzerland.; Universitat Pompeu Fabra, Institute of Evolutionary Biology, Barcelona, Spain.; Passeig de Lluís Companys, Catalan Institution of Research and Advanced Studies, Barcelona, Spain.; Barcelona Institute of Science and Technology, Centre for Genomic Regulation.; Universitat Autònoma de Barcelona, Institut Català de Paleontologia Miquel Crusafont, Barcelona, Spain.; University of Chicago, Department of Medicine, Chicago, IL.

## Abstract

Previously published comparative functional genomic data sets from primates using frozen tissue samples, including many data sets from our own group, were collected and analyzed using non-optimal study designs and analysis approaches. In addition, when samples from multiple tissues were studied in a comparative framework, individual and tissue were confounded. We designed a multi-tissue comparative study of gene expression and DNA methylation in primates that minimizes confounding effects by using a balanced design with respect to species, tissues, and individuals. We also developed a comparative analysis pipeline that minimizes biases due to sequence divergence. We thus present the most comprehensive catalog of similarities and differences in gene expression and methylation levels between livers, kidneys, hearts, and lungs, in humans, chimpanzees, and rhesus macaques. We estimate that overall, only between 7 to 11% (depending on the tissue) of inter-species differences in gene expression levels can be accounted for by corresponding differences in promoter DNA methylation. However, gene expression divergence in conserved tissue-specific genes can be explained by corresponding inter-species methylation changes more often. We end the paper by providing recommendations for effective study design and best practices for meta-data recording for comparative functional genomic studies in primates.

## Introduction

Gene regulatory differences between humans and other primates are hypothesized to underlie human-specific traits [1]. Over the past decade, dozens of comparative genomic studies focused on characterizing mRNA expression level differences between primates in a large number of tissues (e.g., [2-6]), typically focusing on differences between humans and other primates. A few studies have also characterized inter-primate differences in regulatory mechanisms and phenotypes other than gene expression levels, such as DNA methylation levels, chromatin modifications and accessibility, and protein expression levels [7-15].

Comparative functional genomic studies in primates, including from our own lab, often are not designed to test for specific hypotheses. Rather, many of these comparative genome-scale studies aim to build catalogs of similarities and differences in gene regulation between humans and other primates. These catalogs have been shown to be quite useful; for example, they can be used to identify inter-species regulatory changes that have likely evolved under natural selection [3, 4, 6, 13, 16-26], and thereby help us better understand the evolutionary processes that led to adaptations in humans. These catalogs are also used to establish informed models of the relative importance of changes in different molecular mechanisms to regulatory evolution [27, 28]. Comparative catalogs of gene regulation are also used to inform us about ancestral or derived phenotypes that may be relevant to human diseases [29, 30], as well as to help us identify molecular pathways that may be functionally important in the context of complex diseases. Ultimately, comparative catalogs of gene regulatory phenotypes are used to develop and test specific hypotheses regarding the connection between inter-species regulatory changes and physiological, anatomical, and cognitive phenotypic difference between species, in particular between humans and non-human primates.

In primate comparative genomic papers whose main goal is to produce genome-wide catalogs, it can be difficult to detect confounding factors and bias introduced during sample processing or data analysis. Unfortunately, these factors are often neither accounted for nor discussed. In hypothesis driven studies, ignoring confounding effects or introducing bias through errors in study design can lead to erroneous inference and incorrect interpretations. In contrast, in non-hypothesis driven studies, it is typically much more difficult to pinpoint the impact of confounders, because at most, the reported catalog of similarities and differences between species will not be as accurate as it could (and should) have been.

Gene regulatory phenotypes will vary somewhat between studies, regardless of errors in study design, because of differences in the individuals used and the sample processing procedures of each study. Comparative studies in primates, in particular, can typically sample only a small number of individuals from each species, which can lead to incomplete power to detect regulatory differences between species in any given study, and hence to relatively large apparent differences between studies. In light of such expected differences between catalogs derived from different comparative genomic studies, it is difficult to identify specific errors that have resulted from ineffective study design, biases in the analysis, batch effects, or other confounding factors.

As the field of primate comparative genomics has progressed, so too has our understanding of the widespread nature and impact of confounders and other potential biases. For example, earlier comparative studies using gene expression and DNA methylation microarrays, often did not account for the attenuation of hybridization caused by sequence mismatches, which differ between species [6, 17, 18, 31]. Common confounders of the more recent sequence-based comparative studies include individual sampling schemes that are unbalanced across species, and sample processing steps that are segregated by species [2, 10, 21, 32-34]. Systemic differences inherent to the samples, such as differences in material quality between species [2, 35], also remain a concern. In addition, analysis pipelines that do not use orthologous sequences or effective normalization procedures can result in bias [10, 21, 35]. Most comparative genomic studies of humans and non-human primates that we are aware of, including previous studies from our own group [10, 21, 35], suffer from one or more of these weaknesses and caveats.

These considerations are often not explicitly discussed in comparative genomic papers. This puts the onus on the reader to look through the documentation to identify any confounders, and assess their impact on the results and conclusions. Such a task can be quite challenging, as the primate comparative field—and indeed the larger genomics community– lacks consensus regarding meta-data collection and study documentation, particularly around sample and study design reporting.

Another caveat that is shared among all comparative studies in primates is related to difficulty in obtaining multiple tissue samples from the same individual. To date, there have been no published comparative studies in primates that have analyzed multiple tissues sampled from the same individuals across multiple species in a balanced design [30]. As a result, regulatory differences between tissues are always confounded with regulatory differences between individuals. In turn, relative measures of tissue-specific regulatory differences between species are confounded with inter-tissue differences in regulatory variation within species.

It is practically impossible to perform an effective meta-analysis of previous comparative data and account for these caveats and confounders. That said, because inter-species differences typically have large effect sizes, and most comparative studies report general patterns (as we mentioned, specific hypotheses are rarely tested in such studies), we do not expect that accounting for the confounders and batch effects would have a profound impact on the reported results. Specific observations of genes that are reported to be differentially expressed between species may be explained by batch effects or sequence divergence across genes, and the relative divergence of specific genes expressed in different tissues may be affected by the fact that each tissue was sampled from a different individual. The catalogs are not as accurate as they should be; yet, we expect the overall reported patterns to be robust.

Our group and others often use previously published catalogs of comparative data in primates in our different studies. While we do not expect previously observed patterns to be erroneous, we are aware that data on gene-specific inter-species regulatory differences, and especially data that pertain to comparisons of divergence across tissues, may be inaccurate for the reasons we discussed above. We thus designed the current study to produce a new comprehensive catalog of comparative gene expression and DNA methylation data from humans, chimpanzees, and rhesus macaques, attempting to minimize possible confounders. We also developed an analysis pipeline that was designed to minimize bias in comparative studies. We were able to obtain samples from four individuals from each species, sampling four different tissues from each individual. Our study therefore allows us to independently estimate individual, tissue, and species effects on gene expression and DNA methylation levels. We have collected extensive meta-data on all samples, documented practically all sample processing steps, and we provide these data as part of this study. As we subsequently show, even in our carefully designed study, we were not able to exclude all possible partial confounders – this is indeed a difficult challenge when one has to rely on opportunistic sample collection from humans and other apes.

The goal of this study is not to challenge previous conclusions or document specific differences between the current and previous data. Our stated goals are to comparatively study gene regulatory differences, and to estimate the proportion of gene expression variance that can be accounted for by changes in DNA methylation, across tissues and between species. We chose to explore DNA methylation because it is associated with numerous biological processes [36-41] and is hypothesized to impact gene expression levels in mammals [42-44].

We thus aim to provide a new and more accurate comparative catalog of inter-tissue and inter-species differences in gene regulation between humans and other primates, with substantial sample and study design documentation. Overall, we believe that this catalog can be useful for many future applications and can serve as a new benchmark for regulatory divergence in primates.

## Results

### Study design and data collection

To comparatively study gene expression levels and DNA methylation patterns in primates, we collected primary heart, kidney, liver and lung tissue samples from four human, four chimpanzee, and four rhesus macaque individuals (Figure 1A, Additional File 1: Table S1A). From these 48 samples, we harvested RNA and DNA in parallel (Methods). After confirming that the RNA from all samples was of acceptable quality (Additional File 1: Table S1B; Additional File 2: Figure S1A), we performed RNA-sequencing to obtain estimates of gene expression levels. Additional details about the donors, tissue samples, sample processing, and sequencing information can be found in the Methods and Additional File 1: Table S1.

**Figure 1.**
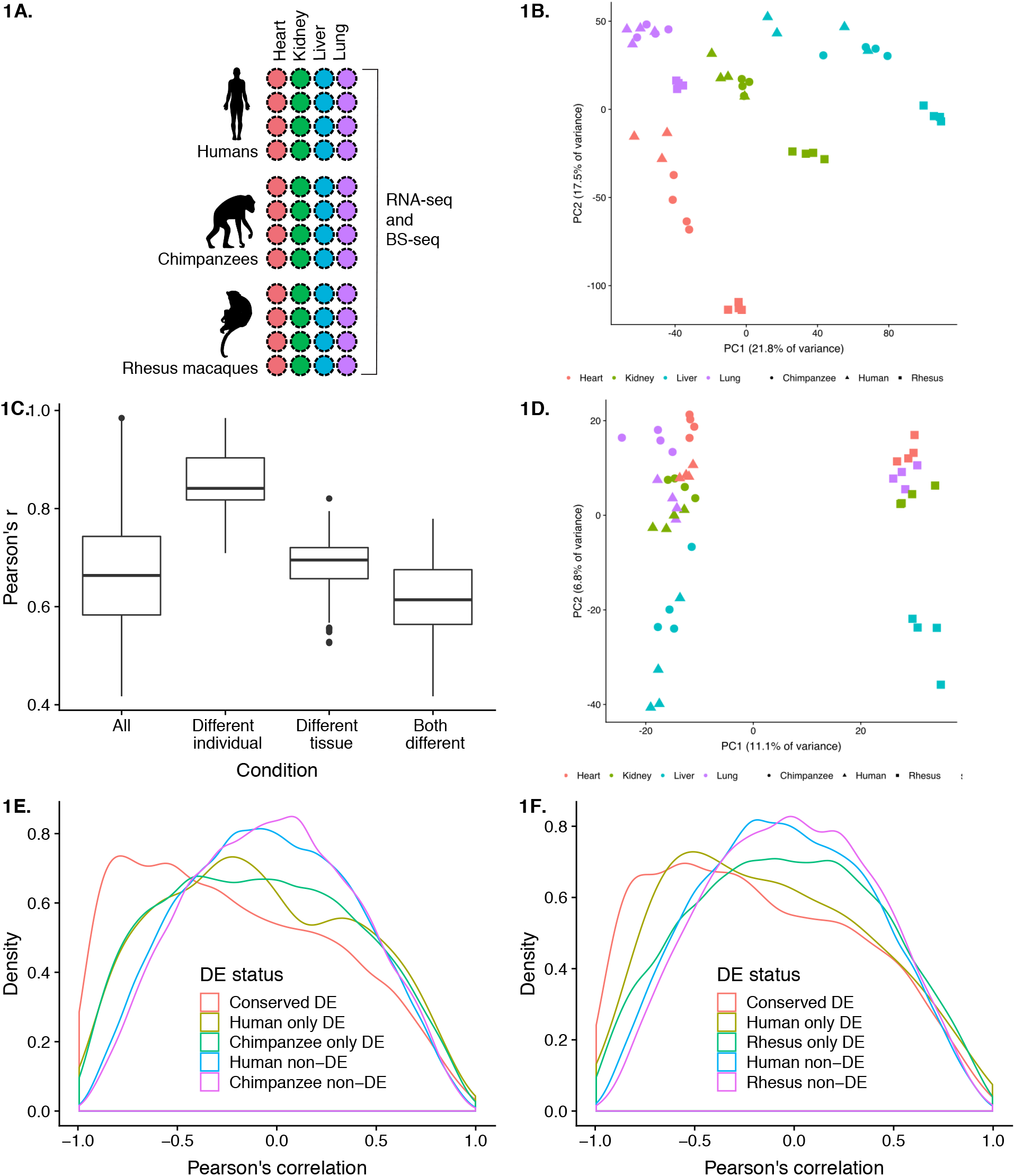
Surveying gene expression and DNA methylation in diverse tissues across primates. (A) Study design. (B) Principal components analysis (PCA) of gene expression levels in 47 samples. (C) Pearson’s correlations of gene expression levels. (D) PCA of average methylation levels 250 bp upsteam and downstream in 47 samples. (E) Density function of the correlation between gene expression and DNA methylation levels in human-chimpanzee orthologous genes. (E) Density function of the correlation between gene expression and DNA methylation levels in genes orthologous across humans and rhesus macaques.

We estimated gene expression levels using an approach designed to prevent biases driven by sequence divergence across the species (similar to the approach of [45]; see Methods). Briefly, we first mapped RNA-sequencing reads to each species’ respective genome, and to compare gene expression levels across species, we only calculated the number of reads mapping to exons that can be classified as clear orthologs across all three species (Additional File 1: Table S1B). We excluded data from genes that were lowly expressed in over half of the samples as well as data from one human heart sample that was an obvious outlier, probably due a sample swap (Additional File 2: Figure S2). We normalized the distribution of gene expression levels to remove systematic expression differences between species (maximizing the number of genes with invariant expression level across species corresponds to our null hypothesis; see Methods). Through this process, we obtained TMM- and cyclic loess-normalized log_2_ counts per million (CPM) values for 12,184 orthologous genes to be used in downstream analyses (Additional File 1: Table S2).

Elements of study design, including sample processing, have previously been shown to impact gene expression data [46]. Consequently, we tested the relationship between a large number of technical factors recorded throughout our experiments and the biological variables of interest in our study, namely tissue and species (Supplementary Information, Additional File 1: Table S3A-B). We found that there were no technical confounders with tissue but two technical factors were confounded with species: time postmortem until collection and RNA extraction date (Additional File 2: Figure 1B-1C). Due to the opportunistic nature of sample collection, these confounders are practically impossible to avoid in comparative studies in primates (especially apes). We discuss possible implications of these confounders throughout the paper.

### Gene expression varies more across tissues than across species

We first examined broad patterns in the gene expression data. A principal component analysis (PCA) indicated that, as expected [2, 47, 48], the primary sources of gene expression variation are tissue (Figure 1B, regression of PC1 by tissue = 0.81; *P* < 10^-14^; regression of PC2 by tissue = 0.70; *P* < 10^-10^; Additional File 1: Tables S1A-B, S3A-B), followed by species (regression of PC2 by species = 0.27; *P* < 10^-3^; Additional File 1: Tables S1A-B, S3A-B). This pattern is also supported by a clustering analysis based on the correlation matrix of pairwise gene expression estimates across samples (Additional File 2: Figure S3). We then confirmed that, globally, gene expression levels across tissues from the same individual are more highly correlated than gene expression levels across tissues from different individuals (Figure 1C). This observation supports the intuitive notion that collecting and analyzing multiple tissues from the same individual is highly desirable in functional genomics studies. Indeed, a rather unusual property of our study design is that we were able to obtain a balanced collection of multiple tissues from the same individuals across multiple primate species.

We next turned our attention to specific gene expression patterns. To analyze the pairwise regulatory differences across tissues and species, we used the framework of a linear model (see Methods). We first identified (at FDR < 1%) 3,695 to 7,027 (depending on the comparison we considered) differences in gene expression levels between tissues, within each species (Table 1, Additional File 1: Table S4). Overall, the patterns of inter-tissue differences in gene expression levels are similar in the three species, significantly more so than expected by chance alone (*P* < 10^-16^, hypergeometric distribution, Methods; Additional File 1: Table S5). A range of 17 – 26% of inter-tissue differentially expressed (DE) genes have conserved inter-tissue expression patterns in all three species (Additional File 1: Table S5). Regardless of species, we found the fewest inter-tissue differentially expressed genes when we considered the contrast between liver and kidney, and the largest number of differentially expressed genes between liver and either heart or lung (Table 1; Table S4B). Unfortunately, since our data were produced from bulk RNA-sequencing, we were unable to determine the impact of cell composition on the number of inter-tissue DE genes.

**Table 1:** Pairwise differentially expressed (DE) genes at FDR 1%.

| **DE between tissues (within species)** | **Heart-Kidney** | **Heart-Liver** | **Heart-Lung** | **Kidney-Liver** | **Kidney-Lung** | **Liver-Lung** |
| --- | --- | --- | --- | --- | --- | --- |
| Human | 4224 | 4776 | 4037 | 4248 | 3695 | 4701 |
| Chimpanzee | 4971 | 6295 | 5365 | 4625 | 4623 | 6247 |
| Rhesus macaque | 5980 | 6933 | 6814 | 5126 | 5721 | 7027 |
| **DE between species** | **Heart** | **Kidney** | **Liver** | **Lung** |  |  |
| Human vs. Chimpanzee | 2195 | 799 | 1364 | 805 |  |  |
| Human vs. Rhesus Macaque | 4098 | 2347 | 2868 | 2833 |  |  |
| Chimpanzee vs. Rhesus Macaque | 2781 | 2286 | 3139 | 1917 |  |  |

We used the same framework of linear modeling to identify gene expression differences between species, within each tissue (Additional File 1: Table S4A). Depending on the tissue and species we considered, we identified between 805 to 4,098 inter-species differentially expressed genes (at FDR = 1%; Table 1). As expected given the known phylogeny of the three species, within each tissue, we classified far fewer differentially expressed genes between humans and chimpanzees than between either of these species and rhesus macaques (Additional File 1: Table S4B).

### Putatively functional tissue-specific gene expression patterns

It is a common notion that genes with tissue-specific expression patterns may underlie tissue-specific functions. Previous catalogs of such patterns in primates were always confounded by the effect of individual variation (because each tissue was sampled from a different individual). To classify tissue-specific genes using our data, we focused on genes that are either up-regulated or down-regulated in a single tissue relative to the other three tissues (within one or more species). We define such genes as having a ‘tissue-specific’ expression pattern, acknowledging that this definition may only be relevant in the context of the four tissues we considered here.

Using this approach and considering the human data across all tissue comparisons, we identified 5,284 genes with tissue-specific expression patterns (FDR 1%, Figure 2). By performing similar analyses using the chimpanzee and rhesus macaque data, we found that the degree of conservation of tissue-specific expression patterns is higher than expected by chance (*P* < 10^-16^ for each tissue; Figure 2). This observation is robust with respect to the statistical cutoffs we used to classify tissue-specific expression patterns (Additional File 1: Table S6), indicating that many of these conserved tissue-specific regulatory patterns are likely of functional significance.

**Figure 2.**
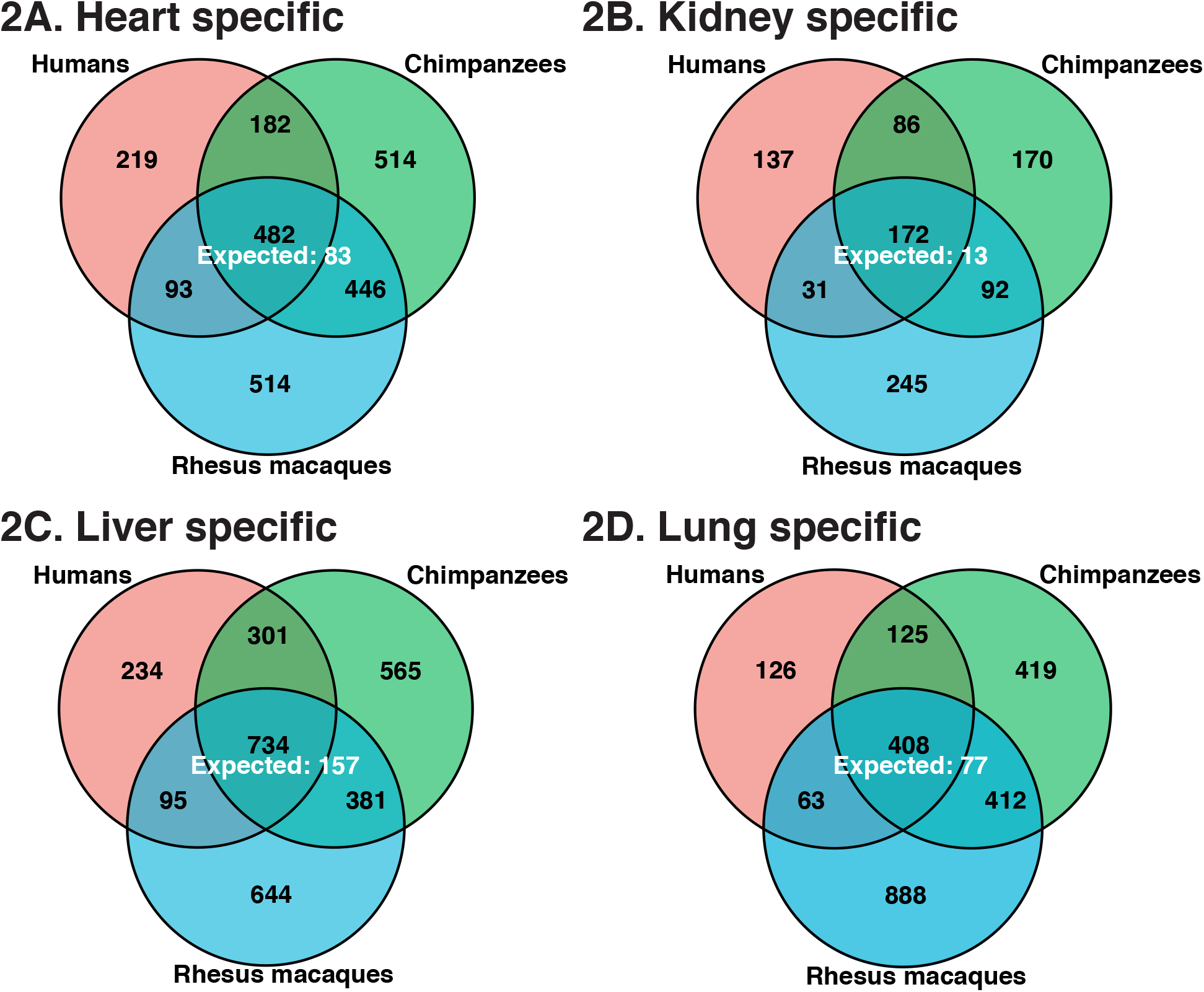
Tissue-specific DE genes (FDR = 0.01) and the expected number of conserved tissue-specific DE genes across all three species, in the (A) heart, (B) kidney, (C) liver, and (D) lung.

We next asked whether the tissue-specific expression patterns we found in a sample of four individuals from each species, are indeed indicative of regulatory patterns in a larger population. To examine this, we considered human GTEx data from the same four tissues we included in our study (see Methods). Because the sample size of the GTEx data is much larger than in our study, the quantity we compared between our data and that from GTEx is the normalized gene expression ranks in the four tissues. For example, in both our and the GTEx data, troponin T2 *(TNNT2)* shows the highest expression in the heart (rank 1) and the lowest expression in the liver (rank 4). Using this approach, we found that 428 (62%) of the 687 genes with a tissue-specific expression pattern exclusively in our human data have the same tissue-based ranked expression in the GTEx data. This observation suggests that tissue-specific expression patterns found in just four individuals are quite often not representative of the regulatory patterns in the larger population. In contrast, however, we found that 1,530 (88%) of the 1,739 genes with a conserved tissue-specific expression pattern based on our data have the same tissue-based ranked expression in the GTEx data. Thus, conserved differential expression significantly increases the confidence of classifying tissue-specific expression patterns in a larger human population (*P* < 10^-16^, difference of proportions test).

To broadly analyze the biological function of genes with conserved tissue-specific expression, we performed a Gene Ontology enrichment analysis (GO, see Methods). We found these genes are indeed highly enriched with functional annotations that are relevant to the corresponding tissue (Additional File 1: Table S7A-D). For example, genes with conserved heart-specific expression patterns were enriched in GO categories related to muscle filament sliding (e.g. *ACTA1, MYL2, MYL6*) and cardiac muscle contraction (e.g. *MYBPC3, MYH7, TNNI3*).

Despite being underpowered to do so (given the small sample size), for the purpose of the catalog, we also identified a small number of genes whose inter-species expression differences across tissues are also tissue-specific. More information about these genes can be found in the Supplementary Information, Methods, and Additional File 1: Table S8.

### Variation in DNA methylation across tissues and species

As we mentioned above, we collected both RNA and DNA from each sample in our study. We used the DNA sample to study DNA methylation patterns through low-coverage whole genome bisulfite sequencing (BS-seq). The bisulfite conversion reaction efficiency was higher than 99.4% for all samples (Additional File 1: Table S1C). Following sequencing, we mapped the high-quality BS-seq reads to *in silico* bisulfite-converted genomes of the corresponding species and removed duplicate reads. We were able to measure methylation level in 12.5M to 22.9M CpG sites per sample, with a minimum coverage of two sequencing reads per site (Additional File 1: Table S1C).

We estimated local methylation levels by smoothing the data across nearby CpG sites (see Methods, Additional File 2: Figure S4); this smoothing procedure has previously been shown to increase the precision of methylation estimates based on low-fold coverage BS-seq data [49]. To facilitate a comparison of methylation levels across species, we identified 10.5M orthologous CpGs in the human and chimpanzee genomes, as well as a smaller set of 2.4M orthologous CpGs in all three primate genomes (see Methods; Additional File 1: Table S1C-E). Unexpectedly, the main sources of variation in the genome-wide methylation data (2.4M orthologous CpG sites) are species, followed by tissue (Additional File 2: Figure S5), as the data from rhesus macaques are quite distinct from the human and chimpanzee data. However, we could not identify a technical explanation for this observation (Supplementary Information, Additional File 1: Table S3C-D). When we focused only on data from CpG sites that are located in or near annotated promoters (see Methods), the similarity of methylation patterns in the same tissues across species was more pronounced, though the major driver of variation is still the separation of rhesus macaques from humans and chimpanzees (Figure 1D). Given the clustering by tissue of the human and chimpanzee data (Additional File 2: Figure S6, Supplementary Information), We believe that the most conservative explanation for the separation of the rhesus data involve technical considerations, although none of our meta-data recorded variables are confounded with species (Methods).

To identify differences in methylation levels between tissues and species we again employed a linear model framework (Methods). Focusing on methylation patterns across tissues within species, we identified between 7,026 to 41,280 differentially methylated regions between tissues, within species (T-DMRs), depending on the pairwise tissue comparisons we considered (Table 2; Additional File 1: Table S9A; [50]). Pairwise comparisons between hearts and lungs showed the lowest number of DMRs, regardless of species (7,026 in rhesus macaques, 8,524 in chimpanzees, 14,208 in humans), while comparisons involving heart and liver showed the largest number of DMRs (22,561 in humans, 28,767 in chimpanzee and 41,280 in rhesus macaques; Table 2). Across all pairwise comparisons, the median length of T-DMRs ranged from 427 bp to 524 bp, with a median of 9-11 CpGs per T-DMR. We found that human T-DMRs overlapped genic and regulatory features significantly more than expected by chance (relative to random, non-T-DMR regions of the same length and CpG density, see Methods). In particular, there is strong enrichment of T-DMRs in CpG island shores [51], 5’UTRs, and active enhancers (as defined by [52]; Additional File 1: Table S9B).

**Table 2:** Pairwise differentially methylated regions (DMRs) in autosomal chromosomes (cutoff recommended by [49])

| **Tissue DMRs (within species)** | **Heart-Kidney** | **Heart-Liver** | **Heart-Lung** | **Kidney-Liver** | **Kidney-Lung** | **Liver-Lung** |
| --- | --- | --- | --- | --- | --- | --- |
| Human | 30291 | 22561 | 14208 | 17910 | 16521 | 12842 |
| Chimpanzee | 17699 | 28767 | 8524 | 23847 | 12076 | 22107 |
| Rhesus macaque | 23023 | 41280 | 7026 | 35910 | 15889 | 32636 |
| **Species DMRs** | **Heart** | **Kidney** | **Liver** | **Lung** |  |  |
| Human vs. Chimpanzee | 14504 | 10667 | 9476 | 8617 |  |  |
| Human vs. Rhesus Macaque | 25539 | 21292 | 21639 | 17696 |  |  |
| Chimpanzee vs. Rhesus Macaque | 19728 | 19585 | 24377 | 15544 |  |  |

We found strong evidence for T-DMR conservation across all three species (*P* < 10^-16^ across all comparisons, at least 50% of bp overlap was required to be considered shared; Additional File 1: Table S10A). Though this level of conservation is higher than expected by chance, we recognize that in each tissue comparison we performed, we had incomplete power to identify T-DMRs and so the true conservation of T-DMR is expected to be even higher. To sidestep this challenge and compare T-DMRs across species more effectively, we considered methylation data from all T-DMR orthologous regions that were classified as such in at least one species. When we performed hierarchical clustering using orthologous DNA methylation data from these T-DMRs, the data clustered first by tissue than by species, even when we included the rhesus macaque data (Additional File 2: Figure S7). This trend is robust with respect to the species used to initially locate T-DMRs (Additional File 2: Figures S8-S9). Thus, our results suggest that in general, inter-tissue methylation differences within a species tend to be conserved, consistent with the observations of previous studies [7, 13, 15, 53, 54].

We next focused specifically on tissue-specific DMRs, as these may contribute to tissue-specific function. In contrast to differences in methylation between any pair of tissues, a tissue-specific DMR is defined as having similar methylation level in three of the tissues we considered, but a significantly different methylation level in the remaining tissue. We found that there were more DMRs specific to liver (3,278 to 11,433 DMRs depending on the species) than to kidney (2,300 to 3,957 DMRs), heart (1,597 to 2,969 DMRs), or lung (453 to 5,018 DMRs, Figure 3; Additional File 1: Table S10B). Tissue-specific DMRs are highly conserved regardless of the comparisons we made (*P* < 10^-13^ for all comparisons, at least 25% bp overlap was required to be considered shared).

**Figure 3.**
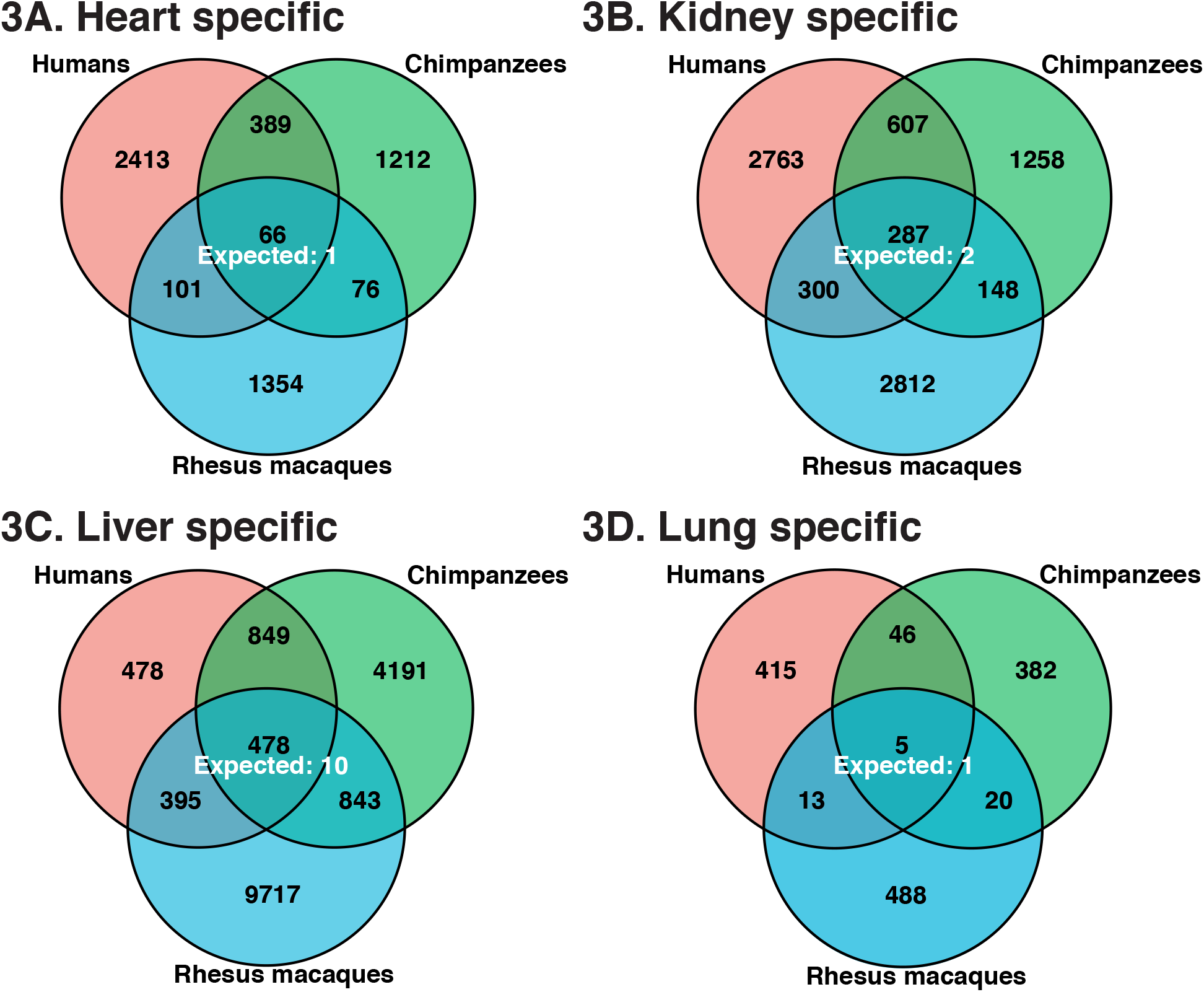
Tissue-specific DMRs (FDR = 0.01) and the expected number of conserved tissue-specific DMRs in the (A) heart, (B) kidney, (C) liver, and (D) lung.

To explore the functional potential of conserved tissue-specific DMRs, we evaluated the overlap between tissue-specific DMRs and genomic regions marked with H3K27ac. Since such genomic regions are often associated with active gene expression [55], we hypothesized that many hypo-methylated tissue-specific DMRs would overlap regions of H3K27ac peaks. Indeed, we found that conserved hypo-methylated tissue-specific DMRs were annotated with this H3K27ac more frequently than tissue-specific DMRs identified only in humans (*P* < 0.001, difference of proportions test; Additional File 1: Table S10C). These observations suggest that conserved tissue-specific DMRs are likely to underlie tissue-specific gene regulation in primates.

### Inter-species differences in gene expression and DNA methylation levels

Our comparative catalog can be used to identify DNA methylation differences that could explain gene expression differences across species and tissues. To do so, we first identified the 7,725 orthologous genes with expression data and corresponding promoter DNA methylation data in humans and chimpanzees, and the 4,155 orthologous genes with the same information for all three species. We then determined to what extent divergence in DNA methylation levels could potentially underlie interspecies differences in gene expression by comparing the gene expression effect size associated with ‘species’ before and after accounting for methylation levels. If methylation patterns underlie inter-species gene expression differences, one might expect the expression effect size associated with ‘species’ to be significantly lower once the corresponding methylation data are taken into account. To determine significant effect size differences, we applied adaptive shrinkage [56] – a flexible Empirical Bayes approach for estimating false discovery rate (Methods).

Considering differentially expressed genes between humans and chimpanzees (in at least one tissue), we found that 11% of genes in the heart, 11% of genes in the lung, 12% of genes in the kidney, and 25% of genes in the liver showed a difference in the effect of species on gene expression levels once average promoter methylation data were accounted for (difference in effect size classified at FSR 5%, Figures 4A and 4E; Additional File 1: Table S11A; Additional File 2: Figure S10; Methods). Our approach was reasonably calibrated, as only a few of the genes showed the unexpected pattern of larger inter-species expression effect after accounting for methylation levels (Figures 4A-4B). As a control analysis, we considered only the genes that were not originally classified as differentially expressed between humans and chimpanzees, and found that the difference in the effect size of species on gene expression levels was reduced in less than 1% of genes once methylation data were accounted for (FSR 5%, Figure 4C, Additional File 1: Table S11A).

**Figure 4.**
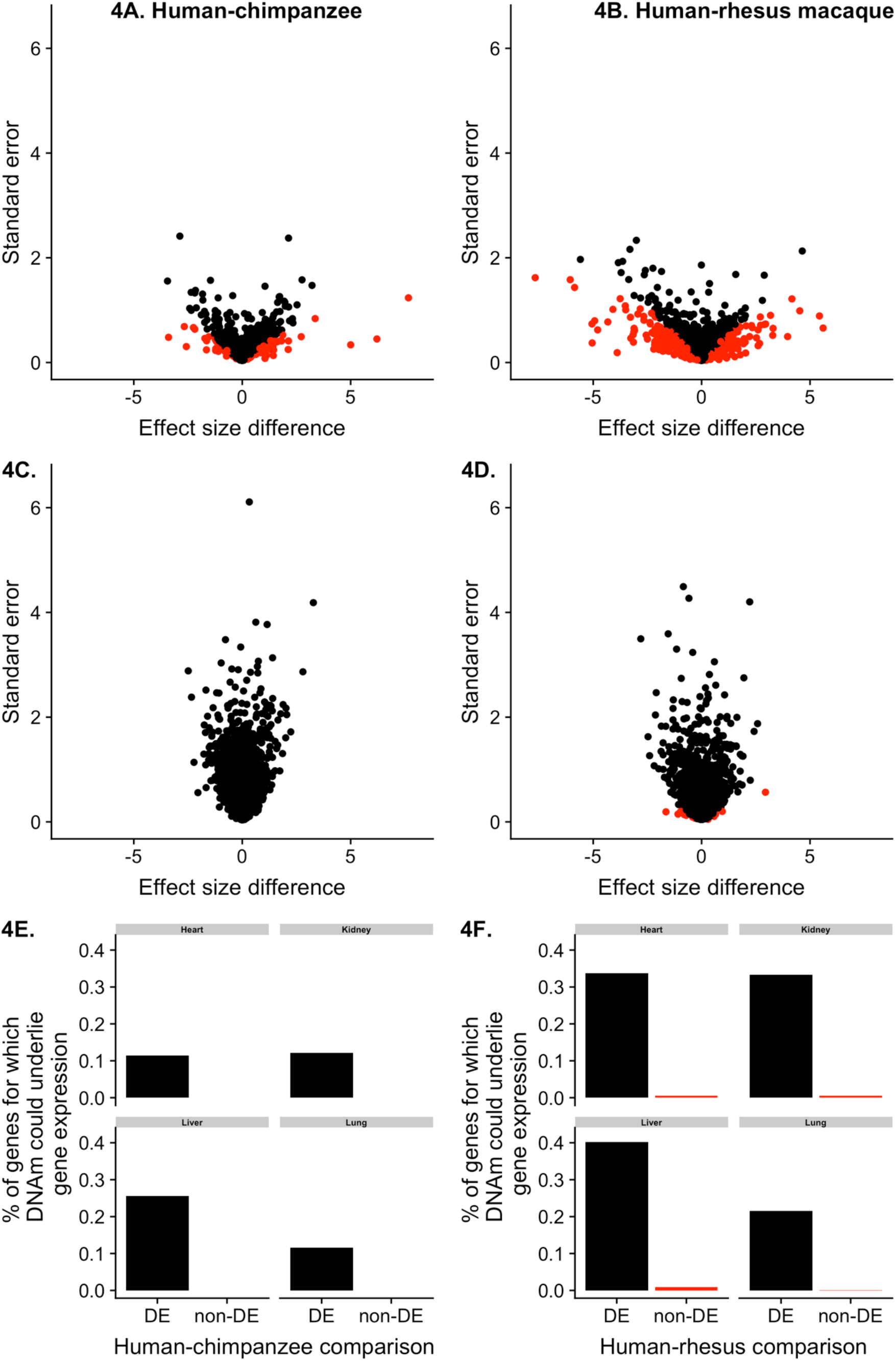
Interspecies DNA methylation and gene expression levels (FDR = 0.05 and FSR = 0.05). **A-D.** Difference in species effect size before and after controlling for DNA methylation levels in genes DE in the (A) human and chimpanzee heart and (B) human and rhesus macaque heart, and non-DE in (C) human and chimpanzee heart and (D) human and rhesus macaque heart. Genes in red are significant at s-value < 0.05. These figures for the other human-chimpanzee tissues and human rhesus macaque tissues can be found in Additional File 2. **E-F.** The percentage of genes for which the evidence for inter-species differences in gene expression levels is reduced after correcting for DNA methylation levels in (E) humans and chimpanzees and in (F) humans and rhesus macaques.

We applied the same approach to the human and rhesus macaque data, and found that the percentage of genes for which gene expression differences could potentially be explained by methylation differences ranges from 21% in the lung to 40% in the liver (DE called at FDR 5% and difference in effect size at FSR 5%, Figure 4B, 4D, and 4F; Additional File 1: Table S11B; Additional File 2: Figure S11). This observation may reflect the more extreme gene expression differences between humans and rhesus macaques than between humans and chimpanzees (prior to accounting for DNA methylation levels, *P* < 0.003 in all tissues, t-test comparing the absolute values of the effect sizes for both groups of DE genes; Figures 4A-4B).

Next, we examined the genes in which DNA methylation differences may underlie inter-tissue gene expression differences. Since our tissue samples were collected from the same individuals, genotype was not confounded with tissue type. Using adaptive shrinkage, we found that 7-25% of inter-tissue gene expression differences could potentially be explained by DNA methylation differences across tissues (FSR 5%; Figures 5A and 5D; Additional File 1: Table S11C-E). When we performed the control analysis and considered only data from genes that were not differentially expressed between tissues, less than 1% of effect sizes differed once we accounted for the methylation data (Figure 5C; Additional File 1: Table S11C-E).

**Figure 5.**
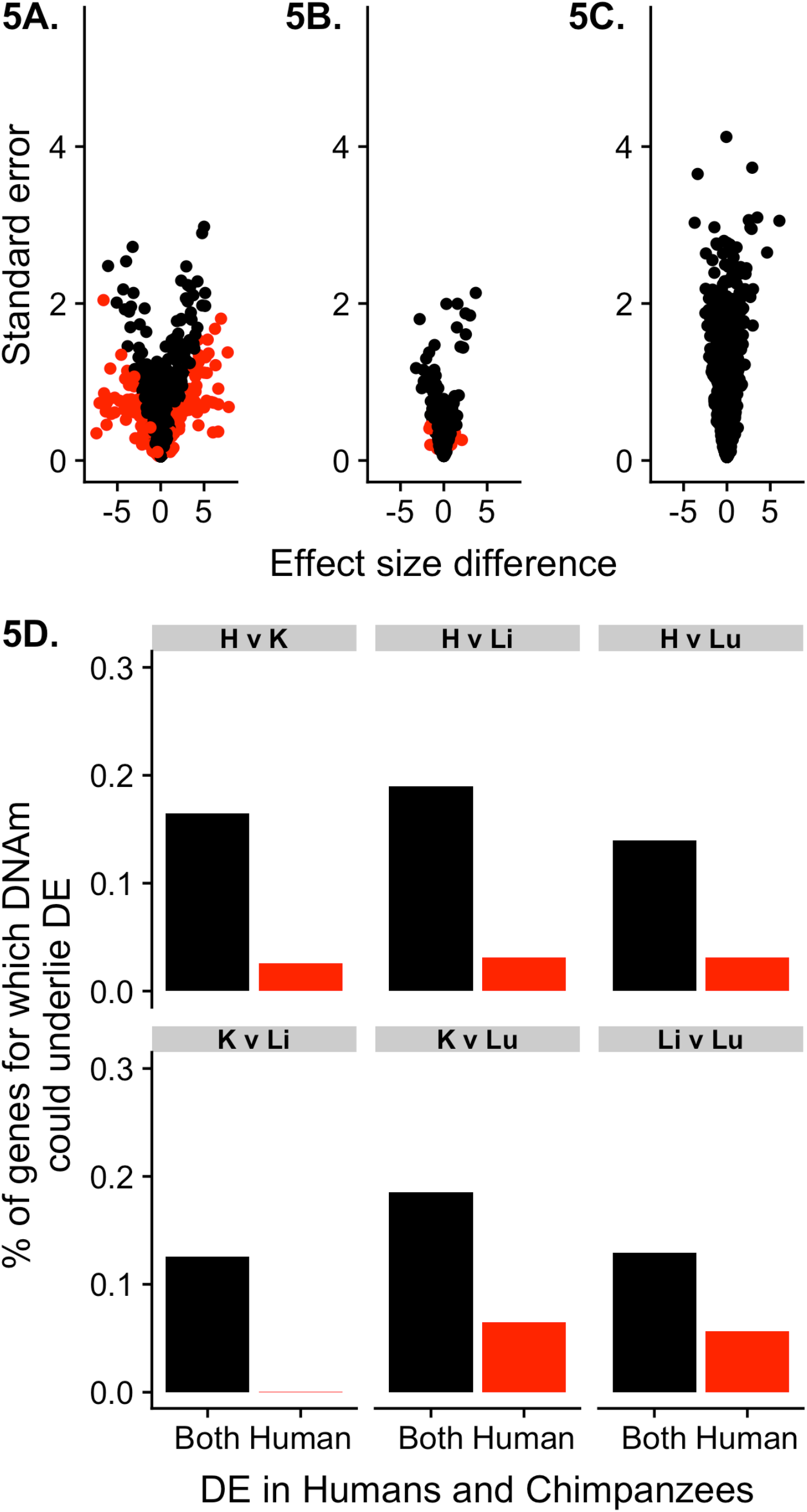
Intertissue DNA methylation and gene expression levels (FDR = 0.05 and FSR = 0.05). **A-C.** Difference in tissue effect size in the human heart versus kidney, before and after controlling for DNA methylation levels. Effect size differences in human heart versus kidney (A) conserved DE genes (DE in humans and chimpanzees), (B) human-specific DE genes (DE in humans but not chimpanzees), and (C) non-DE genes. Genes in red are significant at s-value < 0.05. The same figures for the other human-chimpanzee tissues and human-rhesus macaque tissues can be found in Additional File 2. (D) The percentage of genes for which, after correcting for DNA methylation levels, the evidence for inter-tissue differences in gene expression levels is reduced in human.

Finally, we focused on regulatory patterns that are most likely to be functional; namely, conserved inter-tissue gene regulatory differences. We found that conserved inter-tissue gene expression differences were more likely to be explained by corresponding differences in methylation levels than non-conserved gene expression differences between tissues (minimum difference is 7%, *P* < 0.005 for all comparisons; at FDR < 5% and FSR < 5%; Figure 5A-B, 5D; Additional File 1: Table S11C-E). This observation is robust with respect to the FDR and FSR cutoff used (Additional File 1: Table S11C-E). Indeed, the correlation between methylation and gene expression data is higher for genes with conserved inter-tissue expression patterns compared to genes whose expression patterns were not conserved (Figures 1E-1F).

## Discussion

We designed a comparative study of gene regulation in humans, chimpanzees and rhesus macaques that minimized confounding effects and bias. Because our study was balanced with respect to tissues, individuals, and species, we were able to explore the relationship between DNA methylation levels and gene expression levels at unprecedented resolution. Consistent with previous studies, we found a high degree of conservation in gene expression levels when we considered the same tissue across species [2, 30, 33, 36, 37]. We also found a large number of pairwise tissue DMRs and tissue-specific DMRs to be conserved in the three primates we studied. Our observations are qualitatively consistent with those of previous studies that mostly used microarrays to measure methylation levels [7, 13, 41]. However, the high resolution of our sequence-based DMR data allowed us to examine a much larger number of CpG sites. Thus, we were able to show that while DNA methylation can explain a modest proportion of expression differences between tissues [13], it is more likely to play a role in underlying conserved tissue-specific gene expression levels. While this is an intuitive observation, it has not been explicitly shown previously.

DNA methylation is likely one of the many mechanisms important for early primate gene regulatory evolution [43, 57, 58]. Other mechanisms include sequence difference in regulatory elements, histone modifications, transcription factors, chromatin accessibility, and changes in the 3D genome structure [7-12]. Studies on the impact of these mechanisms on the evolution of gene expression levels are needed to ultimately understand primate gene regulatory evolution. However, performing such functional studies remains difficult due to a reliance on non-renewable frozen tissue samples.

Another limitation of studies that use frozen tissue samples, such as the current study, is that causality can only be inferred. Our inference of casualty in this study is based on the common notion, supported by other studies [36, 37, 42, 44], that methylation levels more often underlie gene expression patterns than vice versa. Based on our data, however, we cannot exclude the possibility that both promoter DNA methylation and gene expression patterns are impacted by a third unobserved mechanism. Our study was not designed to address this or to resolve the causal relationships between different regulatory mechanisms. To do so, we would need a system in which we could directly alter the methylation levels of promoters.

Comparative panels of iPSCs are a renewable system allowing for multiple genomic assays to be performed on the same samples and on multiple derived (differentiated) cell types [59, 60]. We believe that iPSC-derived cell types are useful models for studying spatial and temporal cellular dynamics, especially in humans and apes, where access to other types of material is limited. Unlike frozen tissues, iPSCs are renewable, the differentiation to other cell types can be staged and controlled, and the cell cultures can be perturbed. We acknowledge, however, that most iPSC-derived cell types are better models of fetal rather than adult transcriptional architecture [47-51]. Moreover, tissue samples - even frozen - allow for the study of multiple cell types in a native 3D cellular context [51]. Given the benefits of each type of model system, the field of comparative genomics would benefit from the availability of high-quality data from both iPSC-derived cell types and postmortem tissues.

### Consideration of study design and record keeping

Regardless of the model system used and the types of data that are collected, study design considerations are always critical. Perhaps because comparative studies in primates typically rely on opportunistic sample collection, there are no recognized study design standards that are kept and consistently reported in most existing studies (including many earlier studies from our own group). We thus believe that it is worthwhile to explicitly discuss a few important considerations regarding study design and the recording of meta-data.

Without a balanced study design, it would have been impossible to independently estimate the effects of individual, tissue, and species on our data. Because the sources of confounding factors are difficult to predict in advance, we strongly recommend that samples are collected using a balanced design with respect to as many parameters as possible. These include the distribution of tissue samples per individual, the number of individuals from each species, sex, age range, cause of death and collection time (in the case of post-mortem tissues), or sample collection and cell culturing (in the case of iPSC-based models). All steps of sample processing (RNA extraction, library preparation, etc.) should be done in batches that are randomized or balanced with respect to species, tissue, and any other variables of interest.

Most importantly, all sample processing steps should be recorded in a sample history file that includes anything that happened to any sample. We have documented many of these steps in Additional File 1: Table S1A-E. We recommend, at minimum, recording the following processing steps (in addition to details about the donor individual): sample collection date, RNA extraction batch/date, RIN score, library preparation batch/date, and sequencing details (flowcell, platform, technician, etc.) [32, 52]. This can be an invaluable resource to identify and account for potential confounding factors. Further, these measures may benefit future studies by facilitating effective meta-analysis of multiple data sets, which would help to address the problems of tissue availability and small sample sizes. We believe that, moving forward, it should be a requirement that these meta data are available with every published comparative genomic data set.

## Methods

### Sample Description

We collected heart, kidney (cortex), liver and lung tissues from four individual donors in human *(Homo sapiens*, all of reported Caucasian ethnicity), chimpanzee (*Pan troglodytes*), and Indian rhesus macaque (*Macaca mulatta*), for a total of 48 samples (3 species * 4 tissues * 4 individuals; Figure 1A). The choice of these particular tissues was guided by their relative homogeneity with respect to cellular composition (e.g. [61]), which do not change substantially across primate species. In contrast, other tissues, such as brain subparts, differ substantially in cellular composition across primates [62], which could potentially confound the analyses.

Human samples were obtained from the National Disease Research Interchange (IRB protocol #14378B). Non-human samples were obtained from several sources, including the Yerkes primate center and the Southwest Foundation for Biomedical Research, under IACUC protocol 71619. When possible, samples were collected from adult individuals whose cause of death was unrelated to the tissues studied.

### RNA library preparation and sequencing

In total, we prepared 48 unstranded RNA-sequencing libraries as previously described [63, 64]. Twenty-four barcoded adapters were used to multiplex different samples on two pools of libraries. RNA-sequencing libraries were sequenced on 26 lanes on 4 different flow-cells on an Illumina HiSeq 2500 sequencer in either the Gilad lab or at the University of Chicago Genomics Facility (Additional File 1: Table S1). A relatively small percentage of reads could not be assigned to any sample because their adaptor sequence did not match any of the adaptors used in the study. Therefore, we ran in-house Perl scripts to recover those reads that differed at a single position. Because of a calibration issue with the Gilad lab Illumina HiSeq sequencer, the first and the third flow-cell of the study (16 lanes) yielded a low number of reads. However, this problem did not affect the quality of the reads, so we kept these lanes for the analysis.

### Quantifying the number of RNA-seq reads from orthologous genes

We used FastQC (version 0.10.0; http://www.bioinformatics.bbsrc.ac.uk/projects/fastqc) to generate read quality reports. TrimGalore (version 0.2.8; http://www.bioinformatics.babraham.ac.uk/projects/trim_galore/), a wrapper based on cutadapt (version 1.2.1)[65] was used to trim adaptor sequences from RNA-seq reads. We trimmed using a stringency of 3, and to cut the low-quality ends of reads, using a quality threshold (Phred score) of 20. Reads shorter than 20 nt after trimming were eliminated before mapping (Additional File 1: Table S1).

For each sample, we used TopHat2 (version 2.0.8b) [66] to map the reads to the correct species’ genome: human reads to the hg19 genome and chimpanzee reads to the panTro3 genome. We allowed for up to two mismatches in each read and kept only reads that mapped uniquely. Tophat2 uses unmapped reads to perform gapped alignments to the genome and discover new exon-exon junction sites. For this step, we disabled the coverage-based search, and only 1 mismatch was allowed in the anchor region of the reads (>=8 nt). The minimum intron length was set to 70 nt and the maximum to 50000 nt. This yielded from 28,544,039 (R1H) to 72,808,273 (R4Li) mapped reads across samples (mean = 46,514,692 reads). Mapping rates were between 71% and 94%.

Expression level estimates may be biased across the species due to factors such as mRNA transcript size and different genome annotation qualities. To circumvent these issues, we only retained reads that mapped to a set of 30,030 Ensembl gene orthologous metaexons available for both the 3 genomes, as described and used previously [21, 67, 68]. We defined the number of reads mapped to orthologous genes as the sum of the reads mapping to the orthologous metaexons of each gene. We quantified gene expression levels using the program *coverageBed* from the Bedtools suite.

We performed all downstream analyses in R (versions 3.1.1, 3.2.2, or 3.4.3) unless otherwise stated.

### RNA-seq data transformation and normalization

We calculated the log_2_-transformed counts per million (CPM) from the raw gene counts of each sample using edgeR [69]. We then filtered out lowly expressed genes, keeping only genes with an expression level of log_2_(CPM) > 1.5 in at least 24 of the 48 samples [70]. We normalized the original read counts using the weighted trimmed mean of M-values algorithm (TMM) [70]. This process helped us to account for differences in the read counts at the extremes of the distributions. We then calculated the TMM-normalized log_2_-transformed (CPM) values for each of the genes.

After performing normalization, we performed principal components analysis (PCA) using the TMM-normalized log_2_-transformed (CPM) values of all genes, 1 human heart sample (H1H) clustered with the human livers rather than the hearts. After performing SNP calling on the RNA-seq data (see section below), we found that the SNPs in sample H1H matched those from the other tissues from this individual. We removed this sample (H1H) from the list of the original gene counts. We again filtered for lowly expressed genes keeping only genes with an expression level of log2(CPM) > 1.5 in at least 24 of the 47 samples. We also wanted to allow for small differences in the distributions of gene expression across tissues. Therefore, on the 12,184 remaining genes, we performed a TMM normalization and then performed a cyclic loess normalization with the function *normalizeCyclicLoess* from the R/Bioconductor package limma [71, 72]. To run PCA, we used the R function *prcomp*. For hierarchical clustering, we used unsupervised agglomerative clustering on the correlation matrix of the gene expression data.

In this transformation and normalization process, we were interested in the impact of sample-specific biases in GC content on the gene expression counts. Therefore, we used the WASP pipeline [73] to obtain expected GC-normalized counts. Specifically, we filtered the genes with lowly expressed counts so that only genes with the log_2_-transformed counts per million (CPM) > -5.5 in at least 2 of the 4 samples in each species-tissue pair (e.g. 2/4 chimpanzee hearts) remained. For each of the 16,616 genes that remained, we summed the read depth (raw counts) of the 4 samples in each tissue-species pair. We used the WASP pipeline [73] (https://github.com/bmvdgeijn/WASP/blob/master/CHT/update_total_depth.py) to obtain expected read counts, adjusted for read depth and GC content. The GC content for each of the orthologous metaexons was previously calculated as part of [68]. For each tissue-species pair, the adjusted raw counts and the actual raw counts were highly correlated (> 0.98). Therefore, we did not adjust for read depth or GC content in our RNA-sequencing data.

### SNP calling in the RNA-seq and BS-seq data

We called single nucleotide variants on RNA-seq data from each tissue and sample using standard hard filtering parameters according to GATK recommendations [74]. Briefly, duplicated reads were removed using Picard MarkDuplicates (Rhttp://broadinstitute.github.io/picard). Reads were then subjected to local realignment, base-score recalibration, and candidate-variant calling using the IndelRealigner, TableRecalibration, and HaplotypeCaller tools from GATK [75]. We required a base quality score ≥20. We only considered variants that were observed in at least four of the samples.

We used the same method for the BS-seq data. Through this process, we found that 2 groups of samples had been mislabeled during sequencing: the sample labelled R3Li was actually R2Li, and R3Lu was actually R2Lu.

### Analysis of Technical Variables

To assess whether the study’s biological variables of interest—tissue and species– were confounded with the study’s recorded sample and technical variables, we used an approach described in [45]. We recorded variables related to the samples (e.g. sex), variables specific to gene expression (e.g. RNA-seq flow cell number), and variables related to methylation levels (e.g. number of CpG sites covered) (Additional File 1: Table S1). Briefly, we determined which of our recorded technical variables were significant predictors for each of the gene expression PCs 1-5 using individual linear models for each of the gene expression variables (FDR < 10% for each test). The significant technical variables were then tested against our biological variables of interest, tissue and species, again with individual linear models. For the numerical technical variables, we quantified the strength of these associations using the *P* values from analysis of variance (ANOVA), and used a Chi-squared test (using Monte Carlo simulated *P* values) for the categorical technical variables (significance at FDR < 10%). We repeated the same analysis for methylation data, testing the associations between methylation PCs 1-5 and sample information.

For the 12 RNA-seq related technical variables that were the most highly correlated with tissue or species, we assessed which technical variables constitute the “best set” of independent variables to be included in a linear model for gene expression levels. Because of the partial correlations between the variables, we applied lasso regression using the package “glmnet” [76]. Before performing the analysis, we also protected our variables of interest, tissue and species, in the model for each gene. We summarized each technical variable’s influence across the genes by counting the number of times each technical variable was included in the “best set” of the gene models. We found that none of the technical variables appeared in more than 25% of the best sets (i.e. more than 25% of the gene models). Therefore, we chose not to include these technical variables in our model for testing differential expression.

Finally, during our analysis of technical factors, we discovered that RNA extraction date was confounded with species. In 2012, we extracted RNA from the chimpanzee samples on March 8, from the human samples on three days between March 12-29, and from the rhesus samples on March 6. To test the relationship between the date of RNA extraction and gene expression PCs in humans, we performed individual linear models on PCs 1-5 using RNA extraction date as a predictor. None of the models were statistically significant at FDR 10%, suggesting that tissue type is more highly associated with gene expression levels than RNA extraction date.

### Differential expression analysis using a linear model-based framework

To perform differential expression analysis, we used the same approach as in [45]. We applied a linear model-based empirical Bayes method [77, 78] that accounts for the mean-variance relationship of the RNA-seq read counts, using weights specific to both genes and samples [79]. We implemented this using the R packages *limma* and *voom* [77-79]. This pipeline has previously been shown to perform well with at least 3 samples per condition [80, 81].

We hypothesized that RNA quality may be impacted by postmortem time prior to collection. According to the documentation that we received from the different sites, all of the rhesus macaque samples were collected earlier than all of the chimpanzee samples. These differences could impact RNA quality. Hence, we used RIN score as a proxy for RNA quality, and included RIN score in the linear models.

In the linear models, species, tissue, RIN score, and species-by-tissue interaction terms were modeled as fixed effects. Individual was modeled as a random effect. We used contrast tests in limma to identify genes that were differentially expressed between tissues within each species and across species in the same tissue. We corrected for multiple testing with the Benjamini and Hochberg false discovery rate (FDR) [82]. Genes were considered significantly DE at FDR-adjusted *P* values < 0.01, unless otherwise stated.

To be considered a “tissue-specific DE gene” under our stringent definition, the gene must be in the same direction and statistically significant in all pairwise comparisons including the given tissue but not significant in any comparison without that tissue. For example, for a gene to be classified as having heart-specific upregulation in a given species, the gene needed to be upregulated (a significant, positive effect size) in heart versus liver, heart versus lung, heart versus kidney, but not significantly different between the liver versus lung, liver versus kidney, or kidney versus lung, in the same species. Under the more lenient definition of tissue-specific DE genes, we compared the gene expression level of one tissue to the mean of the other three tissues. To do so, we grouped the three tissues together and again used the limma+voom framework to identify significant differences in one tissue versus the group of the other tissues.

To identify inter-species differences in gene expression patterns across tissues within species (tissue-by-species interactions), we used the limma+voom framework and looked for the significance of tissue-by-group interactions. In one analysis, the groups were Great Ape versus Rhesus macaque and in another analysis, the groups were Human versus non-human primates. To minimize the number of interactions, we compared one tissue relative to a group of the other 3 tissues (e.g. Great Ape versus rhesus macaque heart versus non-heart). Significant tissue-specific interactions were detected using the adaptive shrinkage method, *ashr* [56]. Specifically, for each test, we input the regression estimates from *limma* to *ashr*: regression coefficients, posterior standard errors, and posterior degrees of freedom. We used the default settings in *ashr* to calculate the shrunken regression coefficients (called the “Posterior Mean” in ashr), false discovery rate (FDR or also known as q-value), and false sign rate (FSR or also known as s-value: the probability that sign of the estimated effect size is wrong in either direction). We assigned directionality based on the sign of the posterior mean and determined significance based on the false sign rate.

### Comparing the rank of tissue-specific differentially expressed genes in our dataset to the Genotype-Tissue Expression (GTEx) Project

To benchmark the conserved tissue-specific DE genes, we compared the rank of the gene expression level in our data to the rank of the gene expression level in the GTEx v6 heart, liver, lung, and kidney tissue data [83]. After this comparison, we looked for enrichment of genes with a given rank. To do so, we used the R package *topGO* [84], with the same implementation as in [85]. This implementation included the use of Fisher’s Exact Test, with topGO’s *weight01* algorithm (which takes into account the correlation among GO categories within the graph structure of the program). We then repeated this process for the tissue-specific DE genes identified in humans only.

### Assessing the expected overlap and significance of the observed overlap

We used the process from [13], based on the hypergeometric distribution, to assess the expected overlap of the conserved DE genes and significance of the observed number of conserved DE genes. This process relies on comparing a population proportion to a sample proportion.

We first asked about the overlap of DE genes in humans and chimpanzees. To be conservative, in the case of the human and chimpanzee overlap, we assigned the species with the greater number of genes in the direction of interest as the population and the other species as the sample. To assess the expected overlap in upregulated human and chimpanzee genes in a given tissue, we used the *P* value from the hypergeometric distribution with the following parameters: *m* is the total number of DE genes in the population, *n* is the total number of upregulated genes in a population minus *m*, *q* is the observed overlap of upregulated DE genes (between the humans and chimpanzees), and *k* is the total number of upregulated DE genes in the sample, all within the given tissue. The “expected overlap” is the value at which the maximum likelihood estimate for which *m, n*, and *k* occurs. We then repeated this process for the upregulated and downregulated DE genes, in all 4 tissues separately. To obtain the same statistics for the tissue-specific DE genes, we used only the tissue-specific DE genes, and calculated *n* as the total number of genes upregulated in the tissue of interest compared to the other three tissues in the population minus the number of tissue-specific DE genes in the population.

To calculate these statistics for all 3 species, we used this framework to ask whether the observed overlap between all three species was significant relative to the overlap of the human and chimpanzee DE genes. In the same manner, we used the hypergeometric distribution to assess the expected overlap and significance of the number of conserved tDMRs (tissue differentially methylated regions) and conserved tissue-specific DMRs.

### BS-seq library preparation, sequencing, and mapping

We prepared a total of 48 whole-genome BS-seq libraries from extracted DNA as previously described [86, 87]. To assess the efficiency of the conversion reaction [88], we spiked the extracted DNA with unmethylated lambda phage DNA. For each sample, we prepared at least two libraries, with independent PCR amplifications to minimize PCR duplication rates. The BS-seq libraries were sequenced on 111 lanes on 17 flow-cells on an Illumina HiSeq 2500 sequencer in the Gilad lab or at the University of Chicago Genomics Facility (Additional File 1: Table S1). Reads were single-end and 49 to 59 bp. The distribution of libraries of technical replicates over the flow-cells is described in Additional File 1: Table S1.

Similar to RNA-seq data, we used FastQC to generate quality reports. TrimGalore (version 0.2.8) was used to trim adapter sequences incorporated in the BS-seq reads, using a stringency of 3, and to cut the low-quality ends of reads, using a quality threshold of 20. We eliminated reads shorter than 15 bp post-trimming. We aligned the trimmed reads to the human (hg19, February 2009), chimpanzee (panTro3, October 2010), or rhesus macaque (rheMac2, January 2006) genomes, and to the lambda phage genome using the Bismark aligner (version 0.8.1)[89]. The Bismark aligner maps reads to *in-silico* converted (G to A and C to T) genome sequences using Bowtie (version 1.0.0). This aligner was shown to perform well on benchmark studies [90-92]. We permitted one mismatch in the seed of the alignment, and by default Bismark reports only uniquely mapped reads. Across technical replicates, mapping rates ranged from 49% to 82% (median 76%; Additional File 1: Table S1). We applied the Bismark deduplication script to each technical replicate to remove reads mapped to the same starting genomic position, which likely arise through PCR amplification of the same DNA fragments during library preparation [88]. Across technical replicates, the duplication rates ranged from 2.8% to 44% (median 11%; Additional File 1: Table S1).

We estimated the percentage of methylation at an individual cytosine site by the ratio of the number of cytosines (unconverted) found in mapped reads at this position, to the total number of reads covering this position (sequenced as cytosine or thymine, i.e., converted or unconverted). We calculated methylation estimates genome-wide at CpG sites using the methylation extractor tool from Bismark (version 0.8.1). We additionally collapsed information from both DNA strands (because CpG methylation status is highly symmetrical on opposite strands [93]) to achieve better precision in methylation estimates across the genome. To determine the bisulfite conversion efficiency, we calculated the conversion rate at cytosines from the spiked-in lambda phage DNA (for which coverage ranged from 12× to 107×). We found this rate to be at least 99.4% across technical replicates (Additional File 1: Table S1), increasing confidence in our data.

To obtain CpGs that mapped to multiple species, the chimpanzee and rhesus macaque CpG sites were mapped to human coordinates (hg19) using chain files from ftp://hgdownload.cse.ucsc.edu/goldenPath/hg19/liftOver/ and the LiftOver tool from the UCSC genome browser [94]. These files had previously been filtered for paralogous regions and repeats, but we also removed positions that were not remapped to their original position when we mapped from human back to their original genome. Chimpanzee and rhesus macaque CpG sites were mapped to human, even if their orthologous positions were not a CpG site in human.

After combining all technical replicates, the average genome-wide coverage, calculated at CpG sites shared across the three species, ranged from 1.7× to 5.7× per sample (median of 4×), corresponding to 12M to 23M CpG sites with a coverage of at least 2× (median 19M; Additional File 1: Table S1).

### Methylation level estimate smoothing

Since methylation data from BS-sequencing is typically lower coverage than methylation array data, we first applied a smoothing procedure on raw methylation levels for each sample. It has previously been shown that this procedure increases the precision of low-coverage BS-seq data, and yields methylation estimates that are in excellent agreement with high-coverage BS-seq data without smoothing [49, 95]. To perform the smoothing, we used the BSmooth methold (as implemented in the Bioconductor package *bsseq*, version 0.10.0) [49, 95]– with the default parameters for smoothing– at least 70 CpG sites with methylation data in a smoothing window of at least 1 kb.

After smoothing, we filtered the orthologous CpGs based on coverage to increase confidence in our data. We eliminated sites with > 10x coverage, as the relative sparsity of the data suggests that these reads were likely mapped to repeated regions. For each CpG site in each species/tissue combination, we required an average of 2x coverage and that at least 2 out of 4 individuals had a coverage >= 2x. Post-filtering, we had 2.4M autosomal CpGs orthologous in all three species (10.5 million in humans and chimpanzees only).

### Identifying differentially methylated regions (DMRs)

We were interested in identifying regions exhibiting consistent differences between pairs of tissues or pairs of species, taking biological variation into account. To identify DMRs we used the linear model-based framework in the Bioconductor package bsseq (version 0.10.0)[49]. For a given pairwise comparison (e.g., human liver vs. human heart), the *bsseq* package produces a signal-to-noise statistic for each CpG site similar to a *t*-test statistic, assuming that methylation levels in each condition have equal variance. As recommended by the authors of the package, we used a low-frequency mean correction to improve the marginal distribution of the t-statistics. Similar to previous studies using this methodology, a t-statistics cutoff of –4.6,4.6 was used for significance, [96, 97]. DMRs were defined as regions containing at least three consecutive significant CpGs, an average methylation difference of 10% between conditions, and at least one CpG every 300 bp [49]. We used BEDTools (version 2.26.0) [98] to calculate the number of overlapping DMRs across tissues and/or species [50]. We required overlapping DMRs to have a minimum base pair overlap of at least 25%, unless otherwise stated.

To be considered a tissue-specific DMR, the region was required to be a significant tDMR in 1 tissue compared to the other 3 tissues pairwise (in the same direction) but not a significant tDMR across any of the other 3 tissues in pairwise comparisons. We again used BEDTools to ensure a minimum base pair overlap of at least 25%. Once the tissue-specific DMRs were identified within a species, we then classified them as species-specific or conserved. To be considered conserved (across humans and chimpanzees or all three species), the tissue-specific DMR had to be significant in all species in the comparison and have a minimum base pair overlap of the tDMR of at least 25%.

### Overlap of tissue-specific DMRs with regulatory regions

We extracted the coordinates of the following features from GENCODE annotation (release 19)[99]: exons, first exons, CDS, 3’ and 5’ UTRs, introns, intergenic regions, promoters (-2 kb to +2 kb from TSS [11, 100]) and proximal promoters (-250 nt to +250 nt from TSS [101]) of all genes, or only of protein coding genes. We downloaded coordinates of the CpG islands from the UCSC genome browser [94]. CpG islands (identified as segments of the genome with %G+C > 50%, length > 200 nt, and a ratio of observed over expected number of CG dinucleotides based on number of Gs and Cs in the segment > 0.6 [102]). CpG island shores were defined as 2 kb regions flanking CpG islands, and CpG islands shelves were defined as 2 kb regions outside of CpG islands shores [103]. To test the overlap of tissue-DMRs with enhancers, we used the set of tissue-specific enhancers defined by the FANTOM consortium using CAGE-seq data on primary tissues [52]. To control for the potential effects of CpG density and region length in these analyses, we generated 100 sets of randomly located control regions matching the length and CpG densities of the DMRs in the studied set.

We calculated the overlap of tissue-specific DMRs with H3K27ac in the left ventricle of the heart, kidney, liver, and lung adult tissues from the Epigenome Roadmap [104]. We used the consolidated broad peak data for the left ventricle, liver, and lung (available from http://egg2.wustl.edu/roadmap/data/byFileType/peaks/consolidated/broadPeak/ucsc_compatible/). Since a consolidated version was not available for the kidneys, we used unconsolidated kidney sample numberd 153 (available from http://egg2.wustl.edu/roadmap/data/byFileType/peaks/unconsolidated/broadPeak/ucsc_compatible/). We used BEDTools (version 2.26.0) [98] to calculate the number of tDMRs that overlap an H3K27ac mark.

### Calculating the average methylation levels of conserved promoters

To calculate the DNA methylation levels of orthologous CpGs around the transcription start site (TSS) of orthologous genes, we first had to determine the orthologous TSSs. We began with the 12,184 orthologous genes in our RNA-seq analysis. Of these, we found that 11,131 of these orthologous genes had an hg19 RefSeq TSS annotation (https://sourceforge.net/projects/seqminer/files/Reference%20coordinate/refGene_hg19_TSS.bed/download). We used Lift Over to find orthologous sites in the chimpanzees and rhesus macaque genomes in 9,682 of those 11,131 genes. We then determined which of the hg19 RefSeq TSS annotations were closest to the first hg19 orthologous exon, and repeated this process with the other two species and their respective genomes. We found that 9,604/9,682 of the closest TSS annotations in humans at the same Lift Over coordinates in the other two species. We then calculated the distance between the first orthologous exon to the TSS site in all 3 species individually. To minimize this difference between the 3 species, we filtered all genes with a maximum distance difference across the species of larger than 2500 bp. (For reference, the 75th percentile of the maximum difference in distance was 2,078 bp.) 7,263 autosomal genes remained after this filtering step. 4,155 genes had at least 2 orthologous CpGs 250 bp upstream and 250 bp downstream of the orthologous TSS. We chose a 250 bp window around the TSS based on DNA methylation levels around the promoter in [87] and calculated the average of orthologous CpGs within this window for the 4,155 genes. Using the same method but in humans and chimpanzees only, we found and calculated the average of orthologous CpGs within this window for 7,725 genes.

### Joint analysis of promoter DNA methylation and gene expression levels

To determine whether DNA methylation may underlie interspecies differences in gene expression levels, we used a joint analysis method as described below. For each gene, we analyzed the gene expression levels, along with the accompanying average methylation level 250 bp upstream and downstream of the TSS (found above). For a given tissue, we first determined the effect of species on gene expression levels using a linear model, with species and RIN score as fixed effects (Model 1). Next, we parameterized a linear model attempting to predict expression levels exclusively from methylation levels. We refer to these residuals as “methylation-corrected” gene expression values. We then used these values to again determine the effect of species, this time on gene expression levels “corrected” for methylation, using a linear model with species and RIN score as fixed effects (Model 2). To determine the contribution of DNA methylation levels to inter-species differences in gene expression, we computed the difference in the species effect size between Model 1 and Model 2 for each gene, as well as the standard error of the difference. Large effect size differences between Models 1 and 2 for a given gene suggest that methylation status may be a significant driver of DE. To assess the significance of this difference, we used adaptive shrinkage (ashr) [56] to compute the posterior mean of the differences in the effect sizes, using *vashr*, with the degrees of freedom equal to the number of samples in the linear model minus 2. The shrunken variances from *vashr* were used in the *ashr* posterior mean computation. From this procedure, we obtained the number of genes where species has a significant difference in effect sizes before and after regressing out methylation. We assessed significance using the s-value statistic (false sign rate, FSR [56]). Using the s-values, rather than the q-values, not only takes significance into account but also has the added benefit of assessing our confidence in the direction of the effect.

We performed the above analysis separately for inter-species DE genes and non-DE genes, and in each tissue individually. We identified inter-species DE genes in our tissue of interest as those with a significant species term in the model of species and RIN score as fixed effects. We assessed significance of DE genes at FDR 5%, unless otherwise noted.

We also applied the same analysis framework to determine whether DNA methylation may underlie inter-tissue differences in gene expression levels. For the inter-tissue DE genes and non-DE genes, we replaced “species” with “tissue” as a fixed effect in models 1 and 2. We assessed significance with various FDR and FSR thresholds, as specified in the text.

## Supporting information

## Data Access

Data can be accessed through NCBI’s Gene Expression Omnibus [105] using GEO Series accession number GSE112356 (https://www.ncbi.nlm.nih.gov/geo/query/acc.cgi?acc=GSE112356). Our custom pre-processing script is available upon request to the authors. Pairwise differentially methylated regions can be found at [50]. Data and R scripts used post-processing are available at https://github.com/Lauren-Blake/Reg_Evo_Primates and the results can be viewed at https://lauren-blake.github.io/Regulatory_Evol/analysis/.

## Acknowledgments

Members of the Gilad, Marques-Bonet, Robinson-Rechavi, Stephens, and Pritchard labs provided helpful discussions and comments on the manuscript. In particular, Matthew Stephen provided guidance on integrating the DNA methylation and gene expression data, and Michelle Ward provided helpful comments on a draft of the manuscript. We thank Kasper Hansen, Jenny Tung, and Luis Barreiro for discussions regarding multispecies methylation analysis. We acknowledge Athma Pai for sharing a methylation protocol, Bryce van de Geijn for help with the WASP pipeline, Charlotte Soneson, and Jacob Degner. We also thank the Yerkes Primate Center and Southwest Foundation for Biomedical Research, Anne Stone and Jéssica Hernández Rodríguez for providing and/or helping to ship the tissue samples.

L.E.B. was supported by the National Science Foundation Graduate Research Fellowship (DGE-1144082). Additionally, L.E.B. and I.E. were funded by the Genetics and Gene Regulation Training Grant (T32 GM07197). J.R. was funded by a Swiss NSF postdoc mobility fellowship (PBLAP3-134342) and the Marie Curie International Outgoing Fellowship PRIMATE_REG_EVOL. This project was funded in part by the ORIP/OD P51OD011132 grant. T.M.B. is supported by BFU2017-86471-P (MINECO/FEDER, UE), U01 MH106874 grant, Howard Hughes International Early Career, Obra Social “La Caixa” and Secretaria d’Universitats i Recerca and CERCA Programme del Departament d’Economia i Coneixement de la Generalitat de Catalunya. The content presented in this article is solely the responsibility of the authors and does not necessarily reflect the official views of the funders.

## Additional File 1- Supplementary Tables Legend

**Supplementary Table 1. Recorded variables of interest related to (A) samples, (B) RNA and (C-E) DNA methylation preparation, processing and sequencing.**

**Supplementary Table 2. TMM- and cyclic loess-normalized log_2_ counts per million (CPM) values for 12,184 orthologous genes to be used in downstream analyses.**

**Supplementary Table 3. Technical factor analysis results.** A-B. Results from gene expression levels. (A) Benjamini-Hochberg adjusted P values from from regression models with DNA methylation level PCs 1-5 as response variables and recorded biological and technical variables as explanatory variables. (B) Benjamini-Hochberg adjusted P values from regression models with species and tissue as the response variables and technical factors of interest as the explanatory variables. C-D. Results from global smoothed DNA methylation levels. (C) Benjamini-Hochberg adjusted P values from from regression models with DNA methylation level PCs 1-5 as response variables and recorded biological and technical variables as explanatory variables. (D) Benjamini-Hochberg adjusted P values from regression models with species and tissue as the response variables and technical factors of interest as the explanatory variables.

**Supplementary Table 4. Pairwise inter-tissue and interspecies differential expression statistics.** (A) Differential expression statistics from limma and ash for each pairwise comparison. This includes the Gene name, ENSG gene ID (Gene), log_2_ fold change (logFC), average expression level (AveExpr), t-statistic (t), *P* value (P.value), q-value (adj.P.Val), log-odds (B), an estimate of standard error before adaptive shrinkage (sebetahat), local false sign rate (lfsr), local false discovery rate using estimates from adaptive shrinkage (lfdr), q-value based on the values estimated by adaptive shrinkage (qvalue), s-value based on the values estimated by the adaptive shrinkage (svalue), an adaptive shrinkage-based estimate of beta (beta_est), and an adaptive shrinkage-based estimate of the standard error (se_est) in each pairwise comparison. (B) Number of pairwise inter-tissue and inter-species DE genes at different FDR and false sign rate (FSR) thresholds. FSR is calculated using an adaptive shrinkage method.

**Supplementary Table 5. The most common pattern of inter-tissue gene expression differences is conservation between species is robust with respect to FDR and FSR thresholds.** The observed overlap in the number of pairwise inter-tissue DE genes.

**Supplementary Table 6. Pairwise tissue-specific DE genes are conserved.** The number of tissue-specific genes using a stringent definition (see Methods) across various thresholds.

**Supplementary Table 7. Gene Ontology (GO) results for conserved genes that were upregulated or downregulated only in one tissue (at FDR 1%)**. Results in the (A) heart, (B) kidney, (C) liver, and (D) lung. Within each table, columns A-F represent enriched GO categories that are upregulated or downregulated only in one tissue at a less stringent definition and G-K are at a more stringent definition.

**Supplementary Table 8. Group-by-tissue interactions found in Great Apes and in humans only.** Number of interactions across FDR and FSR thresholds.

**Supplementary Table 9. Summary information for pairwise T-DMRs and S-DMRs** (A) Number of pairwise T-DMRs and S-DMRs (t-statistic > 4.6 or <-4.6). (B) Overlap of T-DMRs with annotated regions, relative to 100 sets of random regions of the same length and CpG density as the T-DMRs.

**Supplementary Table 10. Pairwise T-DMRs and tissue-specific DMRs are conserved.** (A) Conserved pairwise T-DMRs and (B) conserved tissue-specific DMRs.

**Supplementary Table 11. Percentage of genes in which gene expression differences might, at least in part, be explained by DNA methylation levels.** A-B. Percentages for interspecies DE and non-DE genes in (A) humans and chimpanzees and (B) humans and rhesus macaques. C-E. Percentages for inter-tissue conserved DE, non-conserved DE, and non-DE genes in (C) humans, (D) chimpanzees and (E) rhesus macaques.

## Additional File 2- Supplementary Figures Legend

**Supplementary Figure 1. Distributions of potential confounders across biological variables of interest.** (A) RIN score across the samples. (B) RNA extraction date by species (C) PCA of RNA extraction date in humans.

**Supplementary Figure 2. The sample originally labeled as “human 1 heart” is likely a liver from the same human.** (A) One human heart (orange triangle) clusters with the human livers (teal triangles). (B) GATK analysis of the sample labelled “human 1 heart” clusters with the human 1 liver.

**Supplementary Figure 3. Correlation matrix of normalized log2(CPM) gene expression values from 12,184 genes.**

**Supplementary Figure 4. Density function of DNA methylation levels across all species and tissues.** (A) Using raw methylation estimates at the subset of orthologous CpG sites showing a read coverage of at least 5× and no more than 10× in each sample. (B) Using smoothed methylation estimates at all orthologous CpG sites across the three species. (C) Using orthologous CpG sites located in orthologous exons, and excluding sites with C to T SNPs in any of the samples.

**Supplementary Figure 5. Correlation matrix of smoothed DNA methylation levels from all orthologous CpGs.**

**Supplementary Figure 6. Principal components analysis (PCA) in humans and chimpanzee hearts, kidneys, and livers.** (A) Average promoter DNA methylation values. (B) Gene expression levels in the same genes from (A).

**Supplementary Figure 7. When comparing the methylation levels of human T-DMRs and of orthologous regions in the same tissues, clustering is more highly correlated with tissue than species.** Clustering based on human (A) heart-kidney T-DMRs, (B) heart-liver T-DMRs, (C) heart-lung T-DMRs, (D) kidney-liver T-DMRs, (E) kidney-lung T-DMRs, and (F) liver-lung T-DMRs.

**Supplementary Figure 8. When comparing the methylation levels of chimpanzee T-DMRs and of orthologous regions in the same tissues, clustering is more highly correlated with tissue than species.** Clustering based on chimpanzee (A) heart-kidney T-DMRs, (B) heart-liver T-DMRs, (C) heart-lung T-DMRs, (D) kidney-liver T-DMRS, (E) kidney-lung T-DMRS, and (F) liver-lung T-DMRs.

**Supplementary Figure 9. When comparing the methylation levels of rhesus macaque T-DMRs and of orthologous regions in the same tissues, clustering is more highly correlated with tissue than species.** Clustering based on rhesus macaques (A) heart-kidney T-DMRs, (B) heart-liver T-DMRs, (C) heart-lung T-DMRs, (D) kidney-liver T-DMRs, (E) kidney-lung T-DMRs, and (F) liver-lung T-DMRS.

**Supplementary Figure 10. Interspecies DNA methylation and gene expression levels (FDR = 0.05 and FSR = 0.05), in humans and chimpanzees.** Difference in species effect size before and after accounting for DNA methylation levels in genes (A) DE and (B) non-DE in the human and chimpanzee kidney, (C) DE and (D) non-DE in the human and chimpanzee liver, (E) DE and (F) non-DE in the human and chimpanzee lung.

**Supplementary Figure 11. Interspecies DNA methylation and gene expression levels (FDR = 0.05 and FSR = 0.05), in humans and rhesus macaques** Difference in species effect size before and after accounting for DNA methylation levels in genes (A) DE and (B) non-DE in the human and rhesus macaque kidney, (C) DE and (D) non-DE in the human and rhesus macaque liver, (E) DE and (F) non-DE in the human and rhesus macaque lung.

