## Supplementary material for "A comparison of gene expression and DNA methylation patterns across tissues and species"

**Assessing the impact of technical variables on gene expression levels and methylation levels**

We tested the relationship between our technical factors and biological variables of interest, namely tissue and species (see Methods). Through this process, we discovered that RNA extraction date was confounded with species (Additional File 2: Figure S1B). In subsequent analysis with only the human samples (as humans were the only species with samples processed on multiple days), we found that the RNA extraction date did not highly correlate with tissue (Methods, Additional File 2: Figure S1C). Given our particular interest in tissue differences, we do not think that differences in RNA extraction date had a larger impact on variation in gene expression levels than tissue type.

Furthermore, the time of tissue collection post mortem is also confounded with species (Additional File 1: Tables S1A). These differences could impact RNA quality, which can be approximated by RIN score. Indeed, RIN scores were typically higher in rhesus macaques than the other species (Additional File 1: Table S1B; Additional File 2: Figure S1A). As a result, we included RIN score as a covariate when modeling gene expression levels.

The RNA quality may have impacted the number of differentially expressed (DE) genes identified between tissues. This pairwise DE was higher in rhesus macaque than in chimpanzee and human across FDR cutoffs (Additional File 1: Table S5B). This finding is potentially due to the higher sample quality, and therefore lower gene expression level variance, in rhesus macaques.

We also tested for associations between technical factors and biological variables of interest in the BS-seq data. Most of the significant associations were related to DNA methylation levels (e.g. number of orthologous CpGs sites with low methylation, mean methylation level at orthologous CpGs) across species and tissues. We expect the DNA methylation level densities to vary somewhat across tissues [1] and therefore, these inter-tissue differences are likely biological rather than technical.

Additionally, we found a slightly higher overall methylation level in human, compared to chimpanzee and macaque samples (average in humans = 0.664, in chimpanzees = 0.646, in rhesus macaques = 0.626; Additional File 2: Figure S4B), which persisted in the raw, unsmoothed data (Additional File 2: Figure S4A). The distribution of methylation levels could potentially be biased by CpG to TpG homozygous or heterozygous SNPs, which are erroneously inferred as unmethylated CpG sites. If the rate of such SNPs was higher in the studied chimpanzee or macaque individuals compared to human (with respect to their respective reference genome), we could observe differences between species. Since we had performed SNP calling on our RNA-seq dataset, we retained only CpG sites located in orthologous exons, and excluded sites with C to T SNPs in any of the samples. However, we still observed differences between species (Additional File 2: Figure S4C). Moreover, higher methylation rates in humans compared to chimpanzees were previously reported (but rarely discussed) in a diverse set of tissues, using various technologies to measure methylation [2-5]. Therefore, this result may be driven by biases when mapping to different species' genomes.

The coverage on the lambda phage genome was statistically significant (FDR $< 10^{-10}$; Additional File 1: Table S3D). However, further analysis showed that this trend was driven by some low values in the chimpanzee samples, rather than the rhesus macaques (Additional File 1: Table S1D). Indeed, there is no difference in this factor across the human and the rhesus macaque samples ($P = 0.15$, Student's t-test). Therefore, we do not think that coverage on the lambda phage genome can account for the differences in DNA methylation between the Great Apes and the rhesus macaque samples (Figure 1D).

We found evidence for a dependent relationship between species and lane number (Chi squared test, FDR= $10^{-13}$). Since these lanes were spread across multiple flowcells, we do not think that lane substantially contributed to the variance in DNA methylation levels. We also found evidence for a dependent relationship between tissue and library preparation date (Chi squared test, FDR = 0.006). Since the correlation was modest (Pearson's correlation = -0.22) and most tissues within a species had libraries made on multiple days, we chose not to correct for this variable. Finally, we note that sample age has previously been shown to impact methylation status in a subset of genes [6, 7]. DNA methylation levels were weakly positively correlated with age (age quantile relative to the species' average lifespan; Pearson correlation's = 0.18). However, in our factor analysis, this relationship was non-significant (FDR > 10%, Additional File 1: Table S3D). Therefore, we did not correct for this variable.

**Use of adaptive shrinkage and false sign rate to identify tissue-specific genes**

We investigated to what extent our ability to detect tissue-specific genes could be substantially impacted by differences in effect sizes across the species (Additional File 1: Table S6). Therefore, we tested the use of an adaptive shrinkage method [8] to identify genes with a small effect size but consistent direction of effect in each species and used the accompanying false sign rate (FSR) instead of FDR thresholds. The percentage overlap was relatively robust to threshold method (Additional File 1: Table S6). We found that this method increases both the total number of tissue-specific differences and the species-specific differences. However, it also increases the number of conserved tissue-specific gene expression differences in humans and chimpanzees relative to those in chimpanzees and rhesus macaques (Additional File 1: Table S6), more closely reflecting established phylogenetic relationships.

**Identifying inter-species differences between tissues**

Since our data contained multiple tissues and species, we identified genes with interspecies differences between tissues (tissue-by-species interactions). These tissue-by-species interactions are potentially informative for Great Ape evolution (when the contribution of species on gene expression in a given tissue is different between Great Apes and rhesus) and the evolution of human-specific mechanisms in tissues (when the effect of species on gene expression in a given tissue is different between humans and a group containing chimpanzees and rhesus macaques). Using a linear-model based framework, we modeled these differences with tissue-by-species interaction terms (Methods). We found 664 total significant interactions in the Great Ape versus rhesus macaque comparison and 91 in the human versus chimpanzee and rhesus macaque (FDR 1%; Additional File 1: Table S8). Given our sample size and the small effect sizes of these interactions, we are probably underpowered to detect such interactions. To address this, we employed an adaptive shrinkage method [8] to identify genes with a small effect size but consistent direction of effect in each species and used the accompanying false sign rate (FSR) instead of FDR thresholds. This method was used to identify cases where the observed sign of the effect across tissues was different between species. After applying this method, we found 1,006 Great Ape-by-tissue interactions and 257 human-by-tissue interactions (FSR = 1%; Additional File 1: Table S8).

Potentially the most interesting class of tissue-by-species interaction is when species impacts one tissue differently than the other three tissues. Therefore, we used ASH to find 799 Great Ape-by-tissue interactions and 249 human-by-tissue interactions only present in one tissue (FSR = 1%; Additional File 1: Table S8). We defined tissue-by-species specific interactions as interactions with an effect size sign different from the signs of the other interactions (e.g. a positive sign when all other signs are 0 or negative). Unsurprisingly, even after accounting for small effect sizes, there were more tissue-by-species interactions for Great Apes versus rhesus macaques than human-specific ones.

**Promoter DNA methylation quality**

To check our promoter DNA methylation levels in the humans and chimpanzees, we subset the DNA methylation promoter data to the 3 human and chimpanzee tissues tested in a previous study from our lab [1]. Consistent with this previous study, PC1 was more highly correlated with tissue than species and PC2 was more highly correlated with species than tissue (Additional File 2: Figure S6A). Even in this subset of the data, there was more clear separation between tissues in the gene expression levels than the promoter DNA methylation data for these genes (Additional File 2: Figure S6B).

**Identification of differentially methylated regions across species (S-DMRs)**

Using the same method to identify DMRs across tissues, we then identified thousands of DMRs across species (S-DMRs). We found the lowest number of sDMRs on autosomal chromosomes in lungs (8617 DMRs between human and chimpanzees, 17696 DMRs between humans and rhesus macaques, and 15544 between chimpanzees and rhesus macaques) and highest total number in hearts (14504 DMRs between human and chimpanzees, 25539 DMRs between humans and rhesus macaques, and 15544 between chimpanzees and rhesus macaques, Table 2). Similar to the pairwise DE analysis across species, the number of DMRs between species are consistent with known phylogenetic relationships. However, unlike in the pairwise DE analysis across species, the number of S-DMRs is sometimes higher than the number of pairwise T-DMRs. For example, there are more lung S-DMRs than human heart-lung DMRs. This trend is somewhat unexpected given the gene expression data, but consistent with clustering pattern of the methylation data (Figure 1D).
