## Supplementary material for "A comparison of gene expression and DNA methylation patterns across tissues and species"

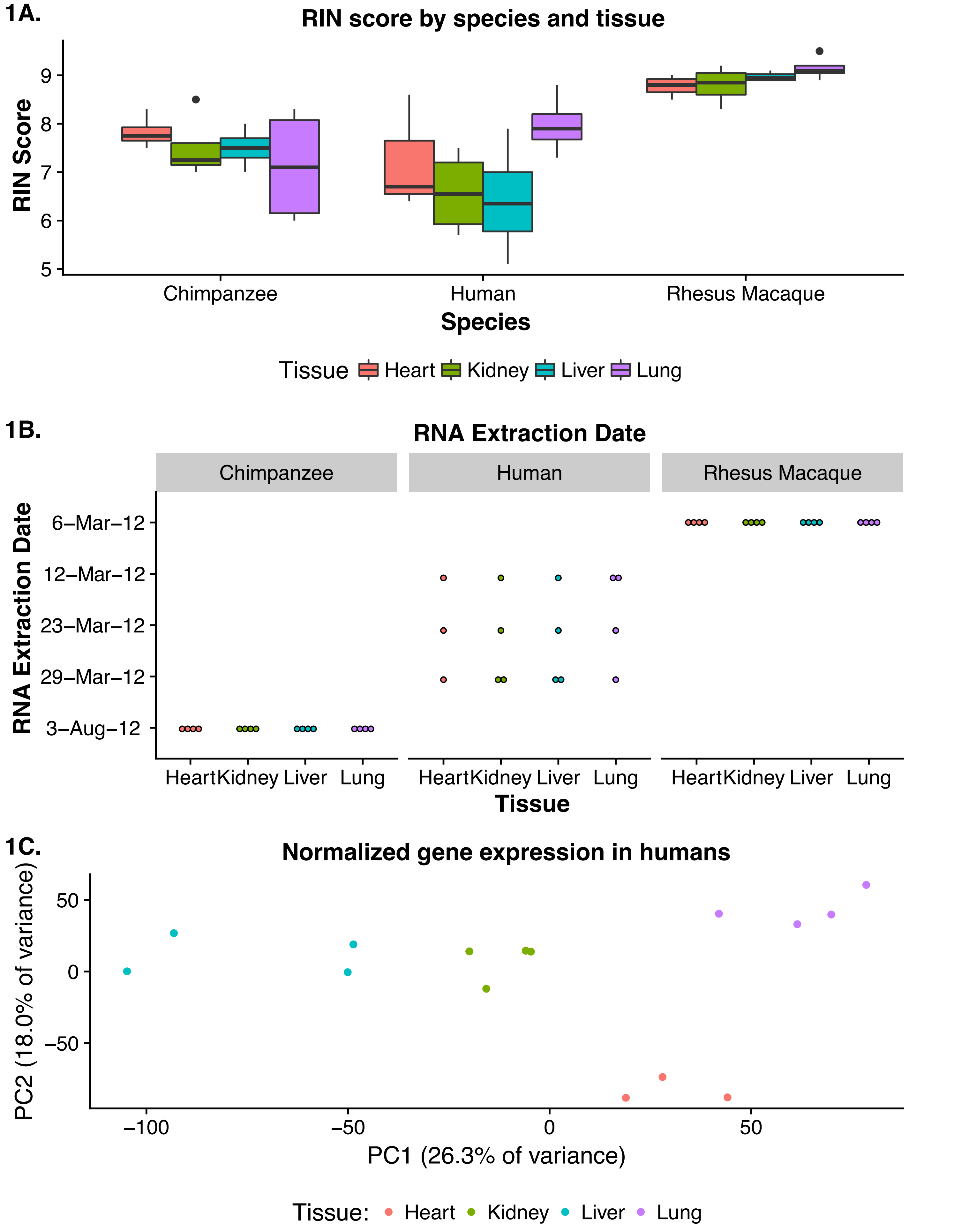
1A.
RIN score by species and tissue
RIN Score
9
8
7
6
5
Chimpanzee
Human
Rhesus Macaque
Species
Tissue
Heart
Kidney
Liver
Lung
1B.
RNA Extraction Date
Chimpanzee
Human
Rhesus Macaque
RNA Extraction Date
6–Mar–12
12–Mar–12
23–Mar–12
29–Mar–12
3–Aug–12
Heart Kidney Liver Lung
Heart Kidney Liver Lung
Heart Kidney Liver Lung
Tissue
1C.
Normalized gene expression in humans
PC2 (18.0% of variance)
50
0
−50
−100
−50
0
50
PC1 (26.3% of variance)
Tissue:
Heart
Kidney
Liver
Lung

2A.

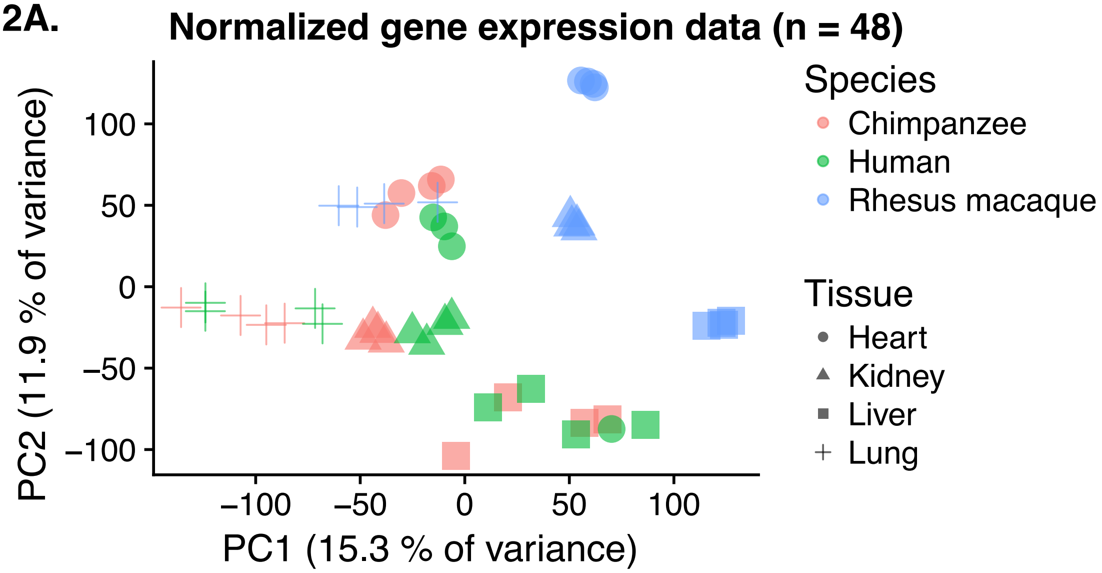
Normalized gene expression data (n = 48)
PC2 (11.9 % of variance)
PC1 (15.3 % of variance)
100
50
0
−50
−100
−100
−50
0
50
100
Species
Chimpanzee
Human
Rhesus macaque
Tissue
Heart
Kidney
Liver
Lung

2B.

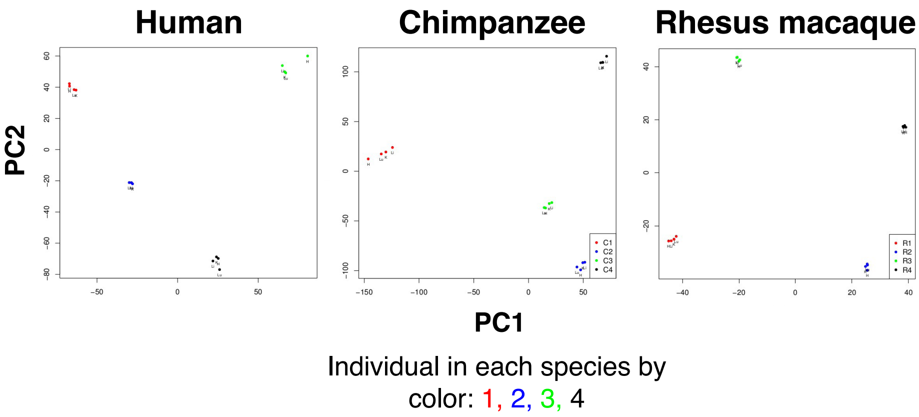
Human
Chimpanzee
Rhesus macaque
PC2
PC1
C1
C2
C3
C4
R1
R2
R3
R4
Individual in each species by
color: 1, 2, 3, 4

3.

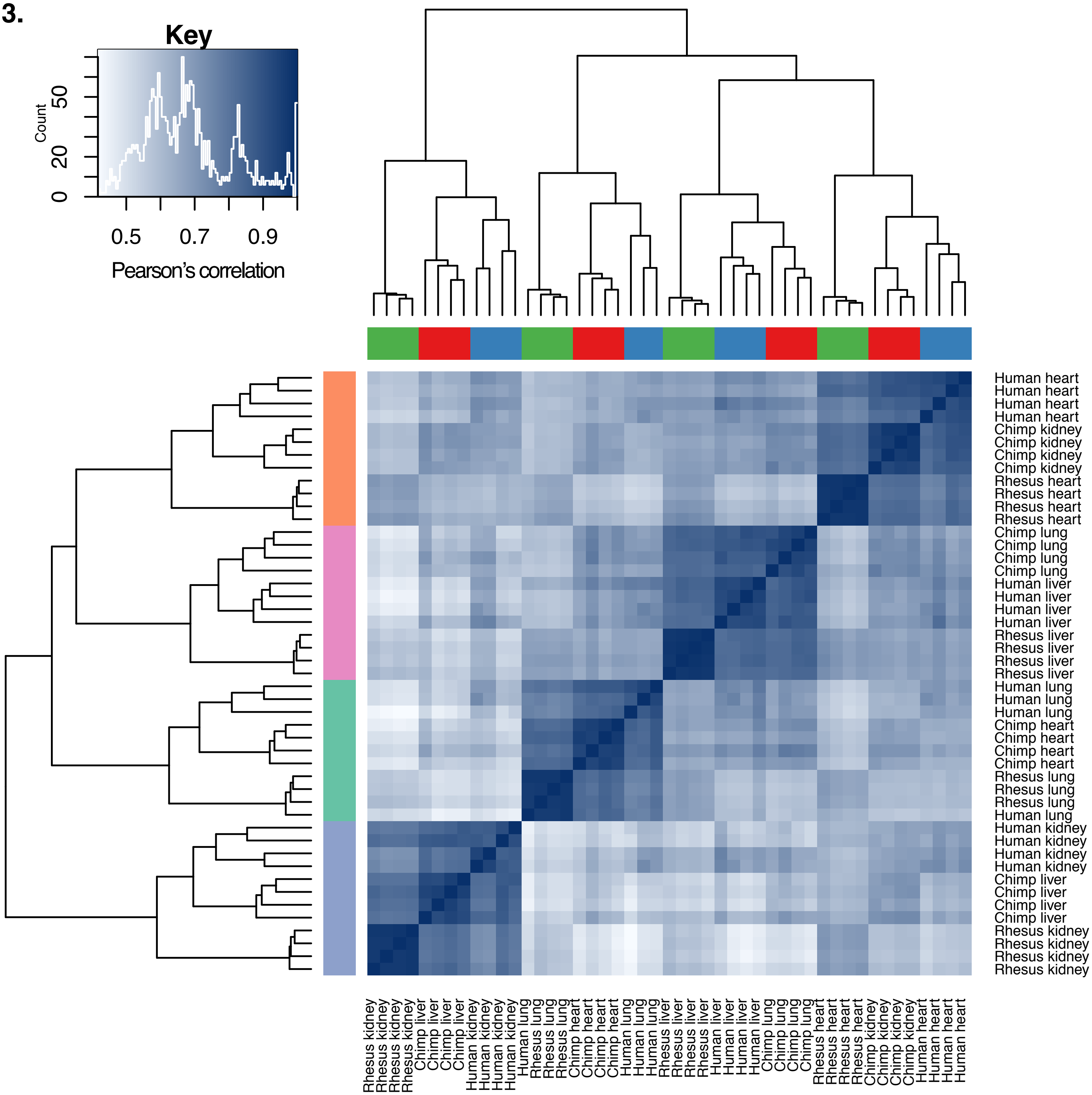
Key
Count
0
20
50
0.5
0.7
0.9
Pearson's correlation
Human heart
Human heart
Human heart
Human heart
Chimp kidney
Chimp kidney
Chimp kidney
Chimp kidney
Rhesus heart
Rhesus heart
Rhesus heart
Rhesus heart
Chimp lung
Chimp lung
Chimp lung
Chimp lung
Human liver
Human liver
Human liver
Human liver
Rhesus liver
Rhesus liver
Rhesus liver
Rhesus liver
Human lung
Human lung
Human lung
Chimp heart
Chimp heart
Chimp heart
Chimp heart
Rhesus lung
Rhesus lung
Rhesus lung
Human lung
Human kidney
Human kidney
Human kidney
Human kidney
Chimp liver
Chimp liver
Chimp liver
Chimp liver
Rhesus kidney
Rhesus kidney
Rhesus kidney
Rhesus kidney

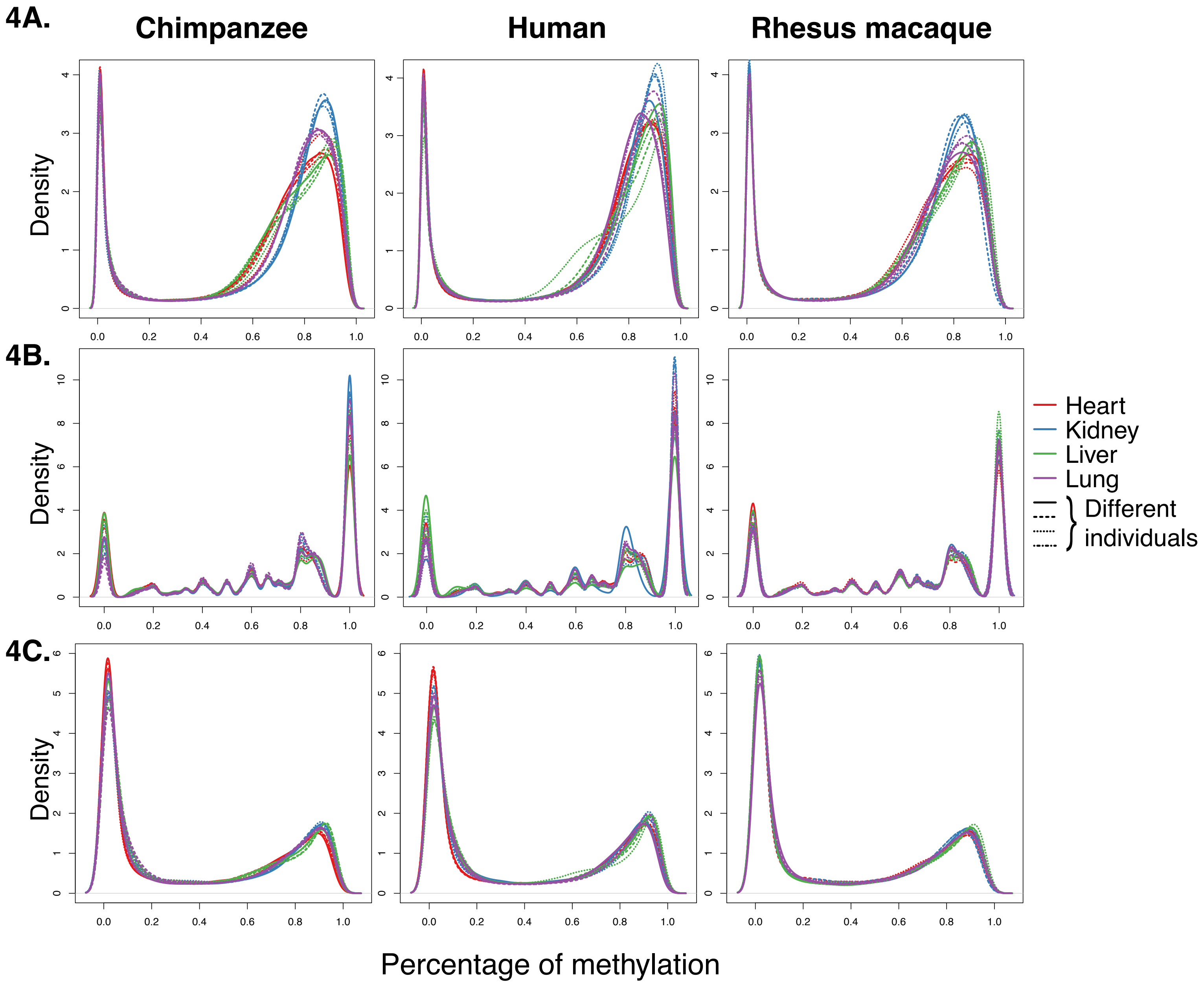

4A.
Chimpanzee
Human
Rhesus macaque
Density
4B.
Density
Heart
Kidney
Liver
Lung
Different individuals
4C.
Density
Percentage of methylation

5.

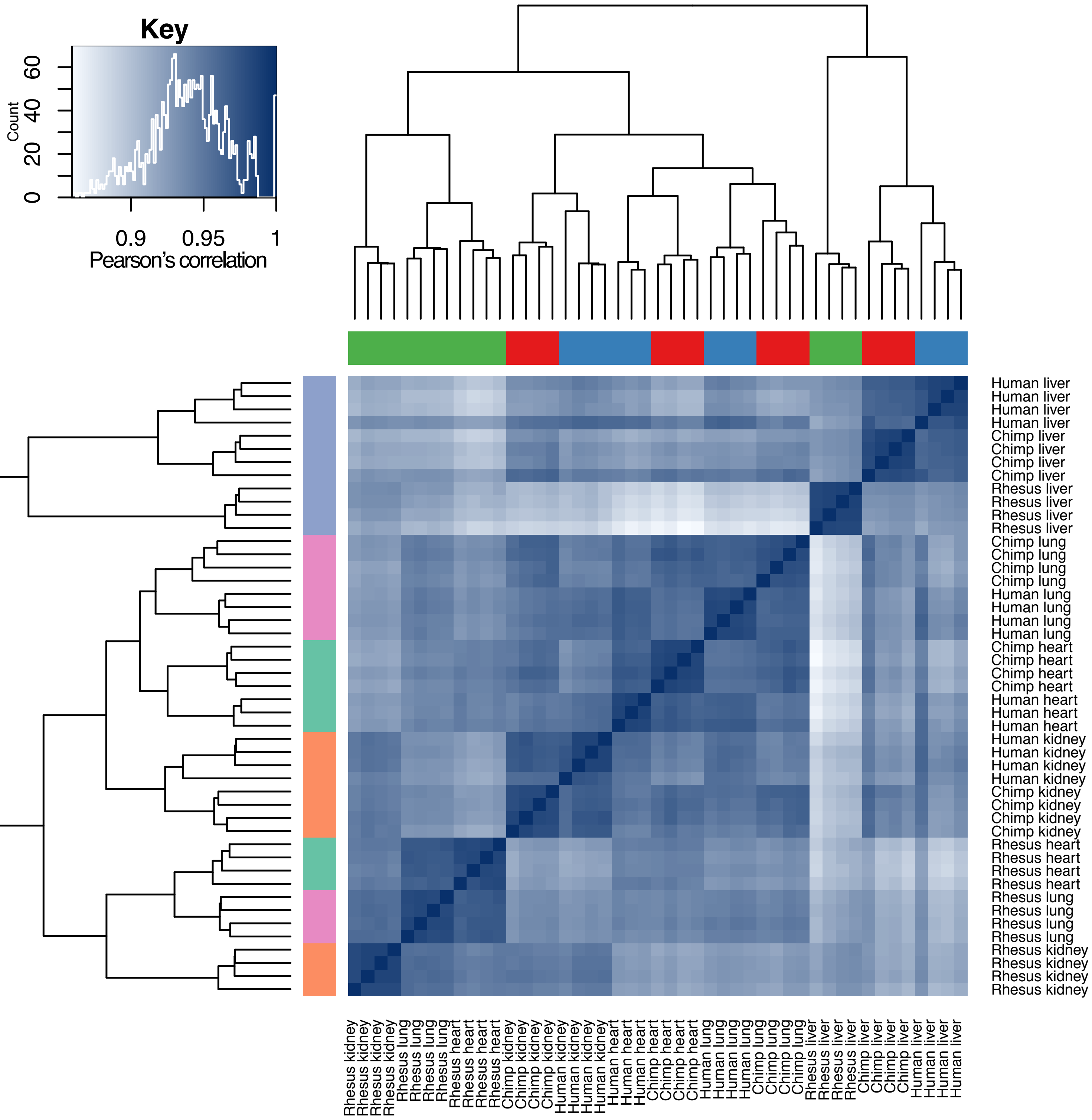

Key
Count
0
20
40
60
0.9
0.95
1
Pearson's correlation
Human liver
Human liver
Human liver
Human liver
Chimp liver
Chimp liver
Chimp liver
Chimp liver
Rhesus liver
Rhesus liver
Rhesus liver
Rhesus liver
Chimp lung
Chimp lung
Chimp lung
Chimp lung
Human lung
Human lung
Human lung
Human lung
Chimp heart
Chimp heart
Chimp heart
Chimp heart
Human heart
Human heart
Human heart
Human kidney
Human kidney
Human kidney
Human kidney
Chimp kidney
Chimp kidney
Chimp kidney
Chimp kidney
Rhesus heart
Rhesus heart
Rhesus heart
Rhesus heart
Rhesus lung
Rhesus lung
Rhesus lung
Rhesus lung
Rhesus kidney
Rhesus kidney
Rhesus kidney
Rhesus kidney

6A.

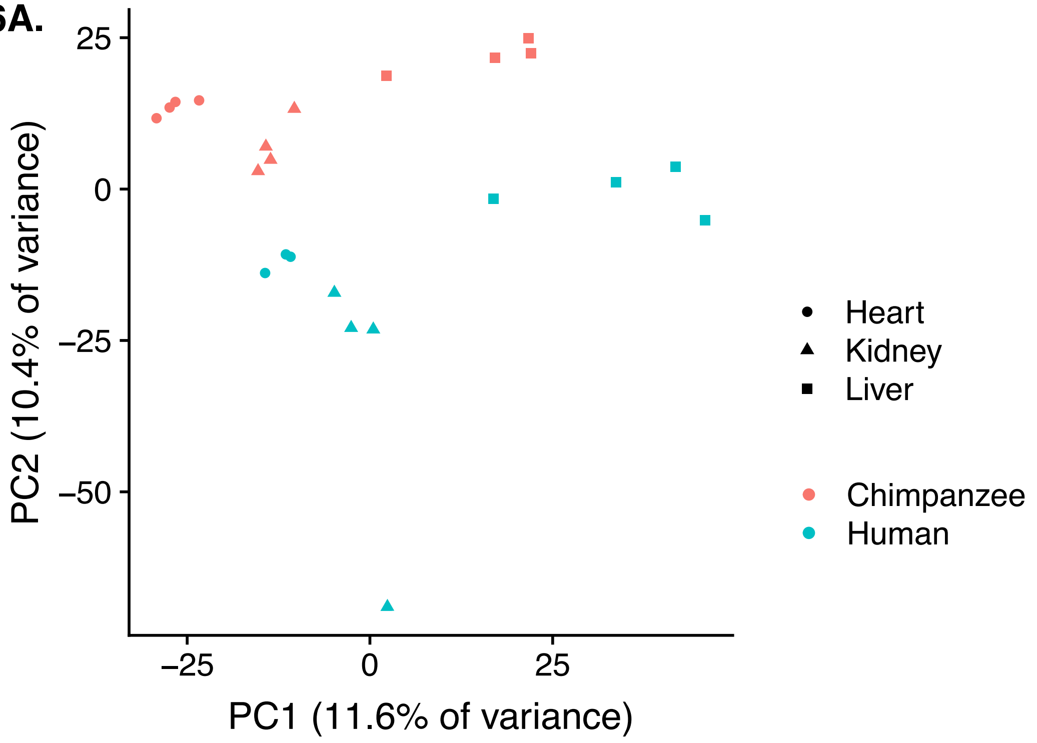
25
0
−25
−50
−25
0
25
PC2 (10.4% of variance)
PC1 (11.6% of variance)
Heart
Kidney
Liver
Chimpanzee
Human

6B.

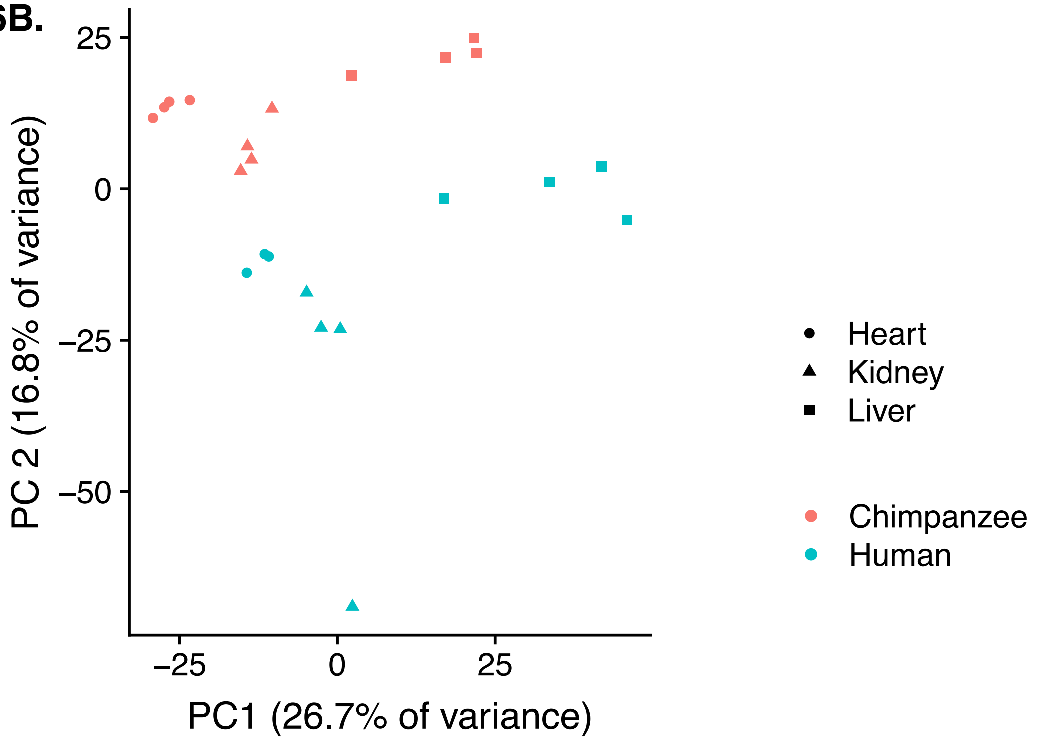
25
0
−25
−50
−25
0
25
PC 2 (16.8% of variance)
PC1 (26.7% of variance)
Heart
Kidney
Liver
Chimpanzee
Human

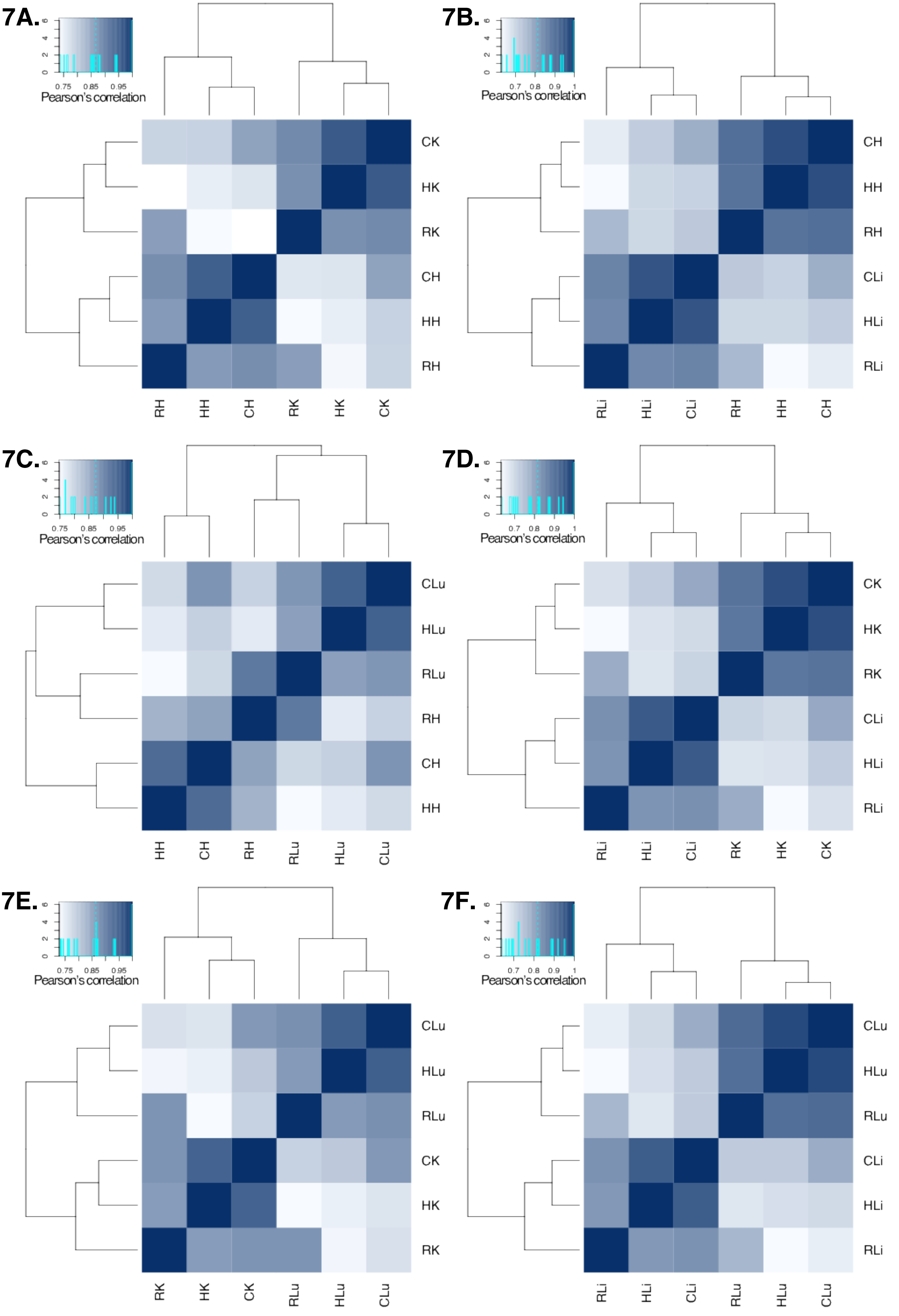
7A.
0
2
4
6
0.75
0.85
0.95
Pearson's correlation
CK
HK
RK
CH
HH
RH
RH
HH
CH
RK
HK
CK
7B.
0
2
4
6
0.7
0.8
0.9
1
Pearson's correlation
CH
HH
RH
CLi
HLi
RLi
RLi
HLi
CLi
RH
HH
CH
7C.
0
2
4
6
0.75
0.85
0.95
Pearson's correlation
CLu
HLu
RLu
RH
CH
HH
HH
CH
RH
RLu
HLu
CLu
7D.
0
2
4
6
0.7
0.8
0.9
1
Pearson's correlation
CK
HK
RK
CLi
HLi
RLi
RLi
HLi
CLi
RK
HK
CK
7E.
0
2
4
6
0.75
0.85
0.95
Pearson's correlation
CLu
HLu
RLu
CK
HK
RK
RK
HK
CK
RLu
HLu
CLu
7F.
0
2
4
6
0.7
0.8
0.9
1
Pearson's correlation
CLu
HLu
RLu
CLi
HLi
RLi
RLi
HLi
CLi
RLu
HLu
CLu

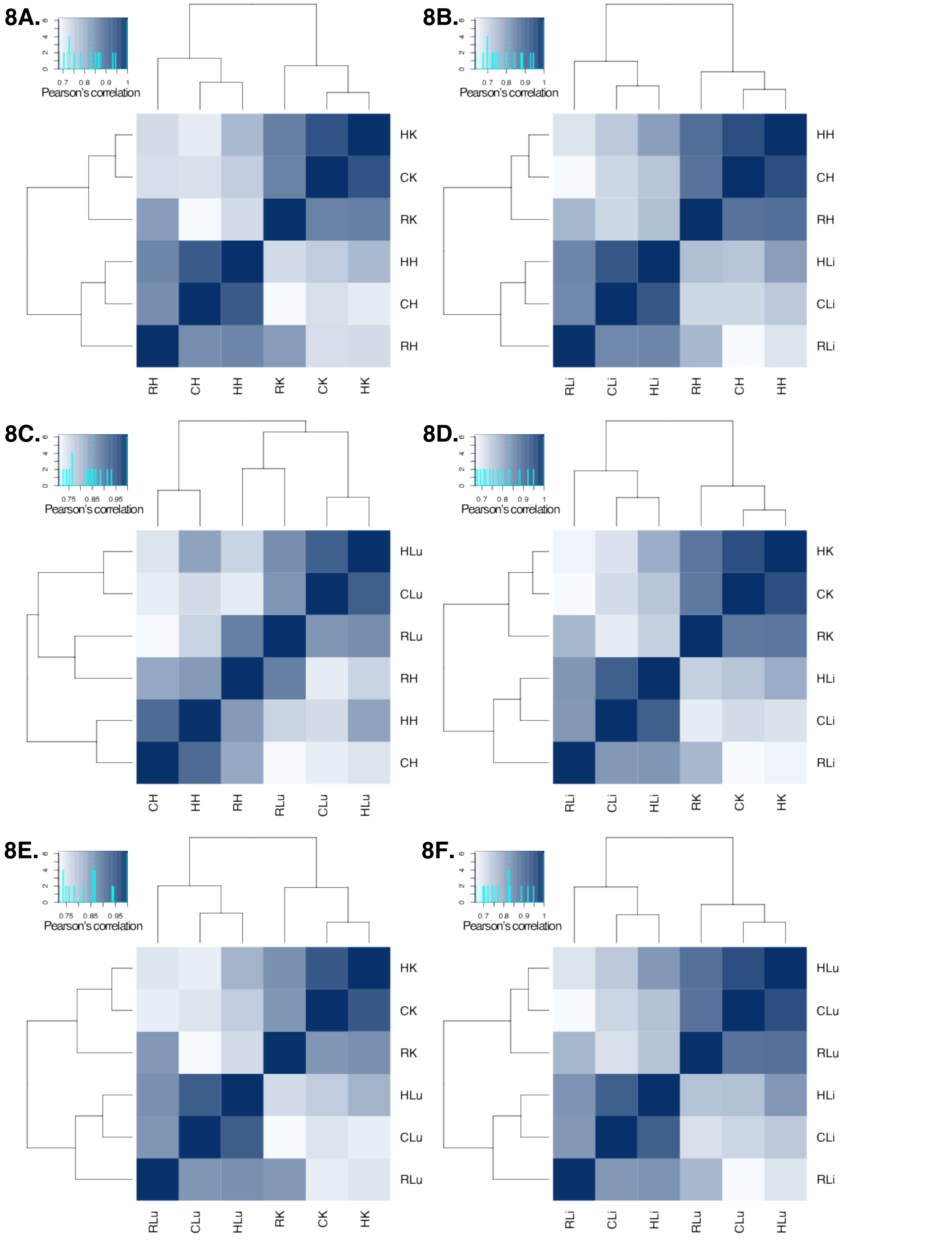

8A.
0 2 4 6
0.7 0.8 0.9 1
Pearson's correlation
HK
CK
RK
HH
CH
RH
RH CH HH RK CK HK
8B.
0 2 4 6
0.7 0.8 0.9 1
Pearson's correlation
HH
CH
RH
HLi
CLi
RLi
RLi CLi HLi RH CH HH
8C.
0 2 4 6
0.75 0.85 0.95
Pearson's correlation
HLu
CLu
RLu
RH
HH
CH
CH HH RH RLu CLu HLu
8D.
0 2 4 6
0.7 0.8 0.9 1
Pearson's correlation
HK
CK
RK
HLi
CLi
RLi
RLi CLi HLi RK CK HK
8E.
0 2 4 6
0.75 0.85 0.95
Pearson's correlation
HK
CK
RK
HLu
CLu
RLu
RLu CLu HLu RK CK HK
8F.
0 2 4 6
0.7 0.8 0.9 1
Pearson's correlation
HLu
CLu
RLu
HLi
CLi
RLi
RLi CLi HLi RLu CLu HLu

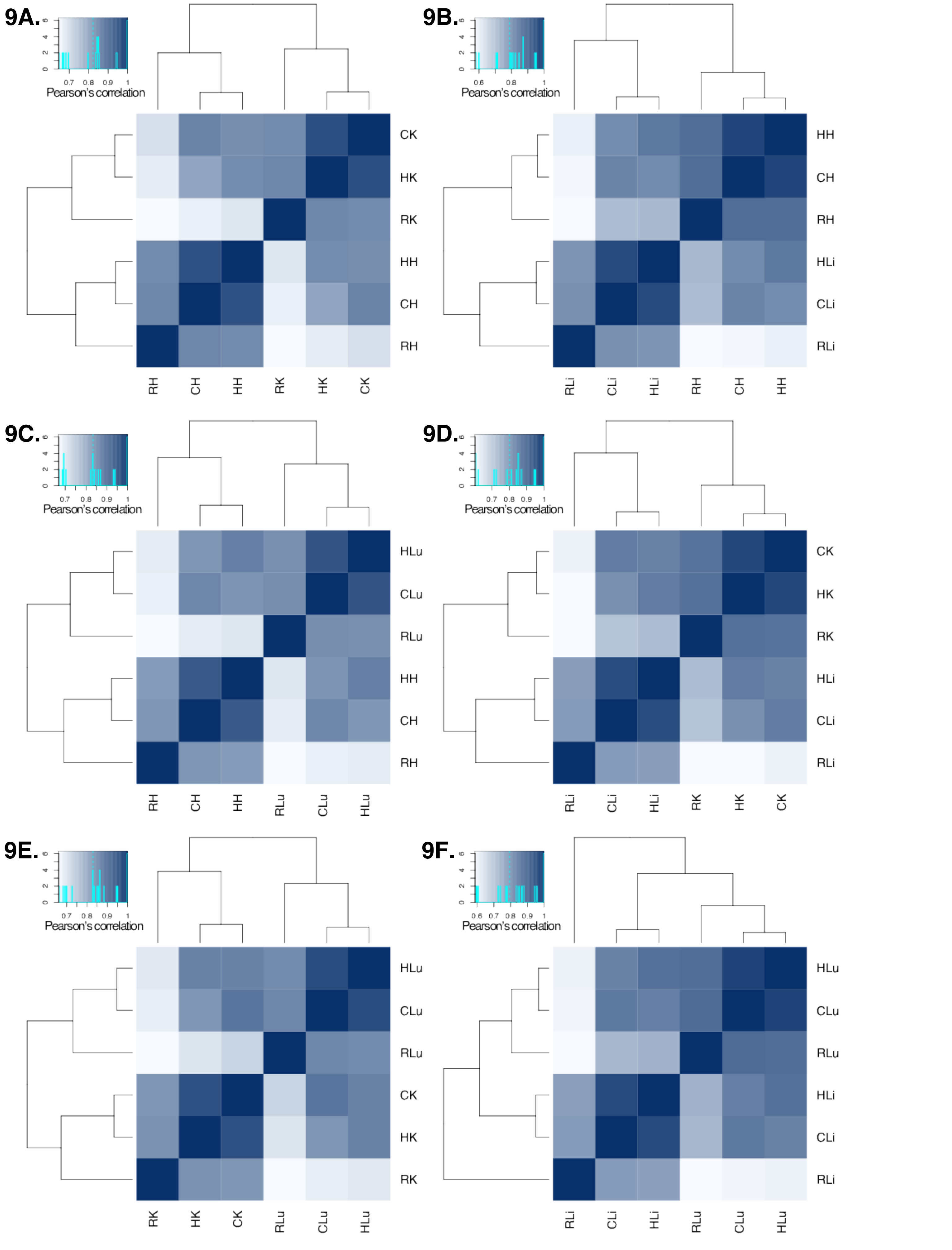
9A.
Pearson's correlation
CK
HK
RK
HH
CH
RH
RH CH HH RK HK CK
9B.
Pearson's correlation
HH
CH
RH
HLi
CLi
RLi
RLi CLi HLi RH CH HH
9C.
Pearson's correlation
HLu
CLu
RLu
HH
CH
RH
RH CH HH RLu CLu HLu
9D.
Pearson's correlation
CK
HK
RK
HLi
CLi
RLi
RLi CLi HLi RK HK CK
9E.
Pearson's correlation
HLu
CLu
RLu
CK
HK
RK
RK HK CK RLu CLu HLu
9F.
Pearson's correlation
HLu
CLu
RLu
HLi
CLi
RLi
RLi CLi HLi RLu CLu HLu

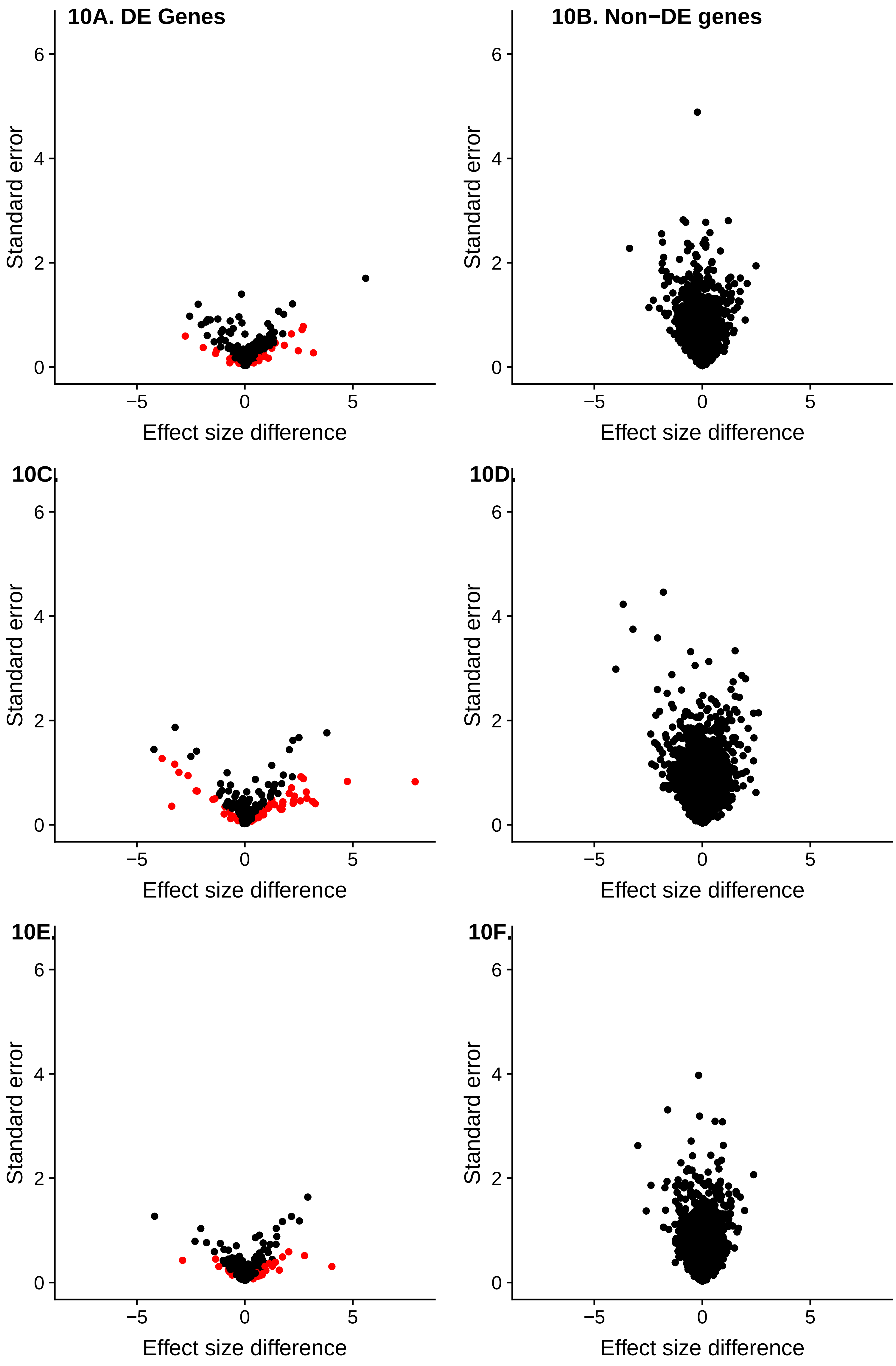
10A. DE Genes
10B. Non−DE genes
10C.
10D.
10E.
10F.
Standard error
Effect size difference
0
2
4
6
−5
0
5

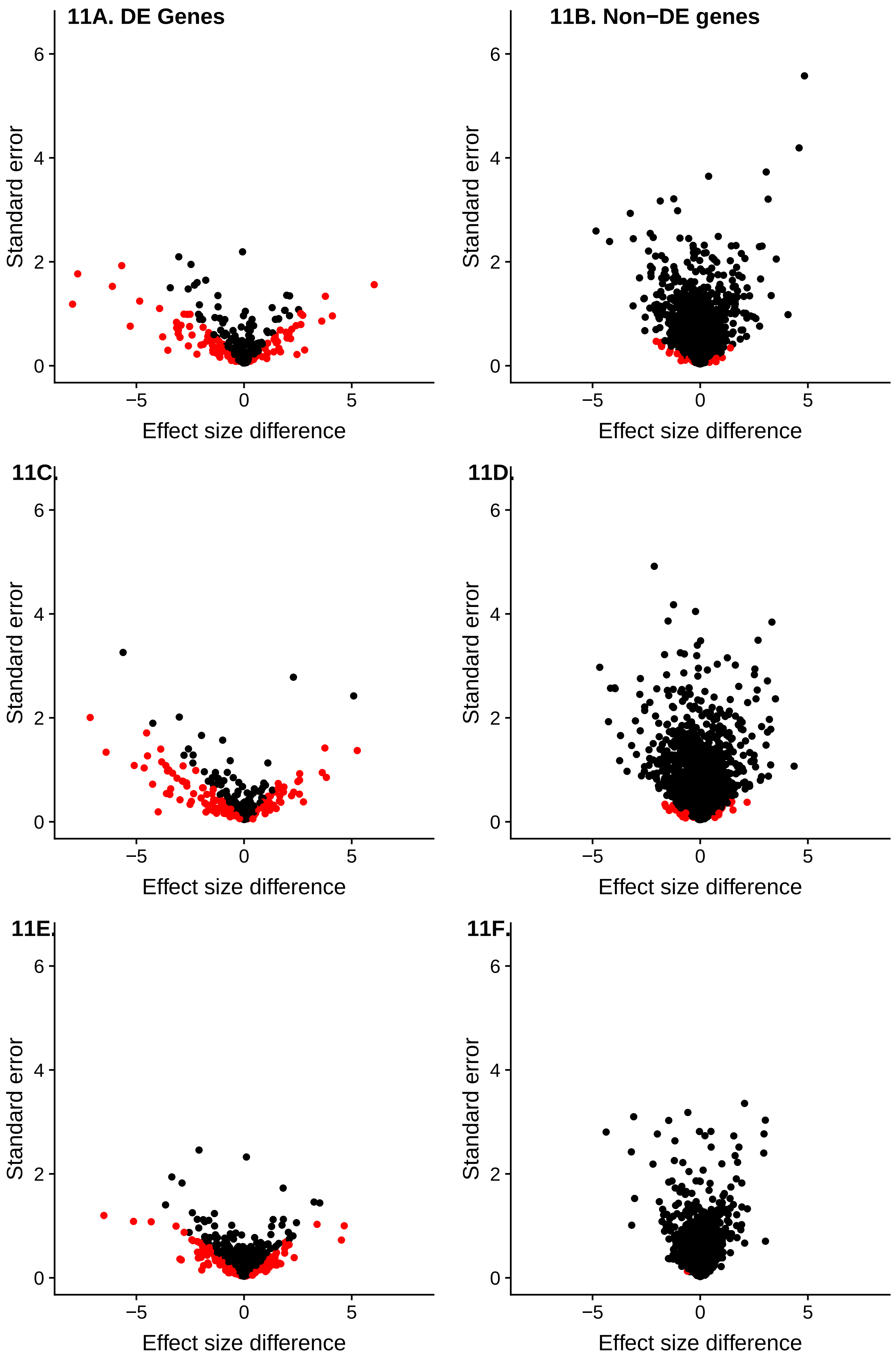
11A. DE Genes
11B. Non−DE genes
11C.
11D.
11E.
11F.
Standard error
Effect size difference
0
2
4
6
−5
0
5
